# Btbd11 regulates glutamatergic synapse organization in GABAergic inhibitory interneurons

**DOI:** 10.64898/2026.06.22.733761

**Authors:** Molly B. Boyer, Shiyu Zhang, Aaron D. Levy, Pascal Schamber, Helene M. Hartman, Thomas A. Blanpied, Alexei M. Bygrave

## Abstract

The mechanisms that underlie glutamatergic synapse organization and function in GABAergic inhibitory interneurons (INs) are not well described, despite evidence that impaired glutamatergic excitation of INs is implicated in psychiatric disorders such as schizophrenia and anxiety. Glutamatergic synapses received by INs have unique basal transmission properties and exhibit distinct synaptic plasticity compared to those received by excitatory neurons, likely due to cell-type specific differences in postsynaptic density (PSD) composition and maintenance mechanisms. In the present study, we show that the interneuron-specific protein Btbd11 regulates excitatory synapse transmission in hippocampal interneurons through promotion of phase separation and support of postsynaptic nanoarchitecture. Btbd11 forms a phase separated protein complex with Psd-95 and TARPγ2 and impacts the stability of TARPγ2 and GluA1 within glutamatergic IN synapses in an expression- and phase separation-dependent manner. Using super resolution imaging, we show that Btbd11 displays nanoscale clustering properties within IN synapses that correlate with Psd-95 nanostructure. Furthermore, genetic deletion of Btbd11 decreases PSD protein expression, reduces synapse size, and disrupts Psd-95 nanocluster organization. These effects manifest as a drastic reduction in glutamatergic synaptic transmission onto INs when Btbd11 is deleted. Together, these data provide insights into a novel cell type-specific synaptic regulatory mechanism in an understudied synapse population.

## Introduction

Glutamatergic synapses in GABAergic inhibitory interneurons (INs) have altered basal transmission, composition, morphology, plasticity, and nanostructure organization^1–7^ compared to excitatory neurons (ENs). The postsynaptic density (PSD) is a highly concentrated assembly of receptors, scaffolding proteins, and signaling molecules that maintains stability over time, yet can be dynamically altered in response to incoming stimuli. The PSD maintains biochemical compartmentalization and stability in INs despite typical localization on the dendritic shaft ^8^ rather than containment within dendritic spines. However, technical limitations in studying the small but diverse IN population has led to a gap in knowledge of cell type-specific molecular mechanisms governing PSD composition and function. Given the link between impaired glutamatergic excitation of INs in psychiatric disease^9–12^, elucidating the unique proteins and mechanisms that govern IN-PSD organization provides an exciting avenue for discovery of potential novel therapeutic targets.

Growing evidence for the PSD as a phase separated condensate provides a mechanism for the stability and maintenance of PSDs, allowing for dynamic molecular exchange in response to incoming signals via glutamatergic transmission while maintaining stability over time^13–15^. Glutamatergic PSD proteins undergo LLPS at physiological conditions, a concentration-dependent process requiring highly specific, multivalent interactions^14,16,17^. Further, LLPS dynamics within the PSD may also provide a mechanism for the formation of nanoclusters (NCs) and trans-synaptic nanodomains through dynamic phase-in-phase segregation ^18–21^. Determining how LLPS is involved in the regulatory mechanisms governing glutamatergic IN-PSD formation and maintenance will provide valuable insight into how INs mediate synaptic signaling, and subsequent disruption in psychiatric disorders.

The dynamic exchange and localization of ionotropic glutamate receptors underlies almost all glutamatergic transmission and plasticity, contributing to memory and cognition. Synaptic AMPAR clustering depends critically on membrane-associated guanylate kinases (MAGUKs), the major scaffolding family at the PSD, which organize receptors through interactions with auxiliary subunits such as transmembrane AMPAR regulatory proteins (TARPs) ^22–25^. While some interactions between PSD signaling molecules are well-characterized in ENs, such as the Psd-95/TARP/AMPAR complex ^22,26,26–31^, the mechanisms of PSD complex regulation underlying synapse organization are not fully understood, especially in INs.

Btbd11 has recently been identified as an IN-specific, glutamatergic synapse-enriched protein that supports synaptic and circuit function ^32^. Btbd11 has been shown to bind directly to and undergo LLPS with Psd-95 ^32^, which could contribute to the stability and function of IN-PSD. Structurally, Btbd11 has a PDZ-binding motif (PBM) that facilitates the interaction with Psd-95, ankyrin repeats, and a disordered N-terminal region, reminiscent of other known PSD components that can undergo LLPS ^17,33^. Ablation of Btbd11 disrupts PV-IN function, synapse size, and circuit activity, suggesting postsynaptic disruptions.

Here, we use biochemistry, live-cell imaging and a combination of confocal and super resolution microscopy to examine Btbd11 protein interactions, complex formation, mobility, expression, and the relevance of phase separation in INs. We show Btbd11 forms a protein complex with Psd-95 and key glutamatergic receptor-associated proteins essential for AMPA receptor synapse targeting. Further, Btbd11 impacts the stability of key glutamatergic signaling proteins PSD-95, TARPγ2, and GluA1 in a phase separation-dependent manner. We also show that Btbd11’s nanoscale distribution is related to Psd-95, and that loss of Btbd11 diminishes Psd-95 NC size and number. Finally, we observe that Btbd11 ablation leads to a decrease in spontaneous postsynaptic calcium transients. Together, these data identify a novel cell type-specific synaptic regulatory mechanism governing IN glutamatergic synapse function and provide insight into the role of phase separation at the synapse.

## Results

### Fine-mapping the Btbd11-Psd-95 interaction and protein complex formation with TARPγ2

We previously found that Btbd11 interacts directly with Psd-95 to stabilize its localization at glutamatergic synapses within INs, an interaction occurring via Btbd11’s C-terminal PDZ-binding motif (PBM) binding to Psd-95’s PDZ1/PDZ2 domains ^32^. To determine which of Psd-95’s PDZ domains binds to Btbd11, we used predictive modeling machine learning software AlphaFold3 ^34^ to model the interaction between Btbd11 and Psd-95. The software predicted that Btbd11 preferentially and specifically binds to the PDZ2 domain (Figure 1A, Supplemental Figure 1A). To experimentally verify PDZ2 as the primary interaction domain, we co-expressed GST-Btbd11 with full-length or Psd-95-mCherry PDZ mutants in HEK cells to perform GST immunoprecipitations (IPs). PDZ mutants comprised a six-residue deletion in the βB/βC loop of PDZ1, PDZ2, or both ^35^. Western blots of the lysates determined that compared to full-length Psd-95, mutation of the PDZ1 domain reduced Btbd11-Psd-95 binding while mutations of either PDZ2 alone or PDZ1/2 together completely abolished Btbd11-Psd-95 binding (Figure 1B-C).

**Figure 1.**
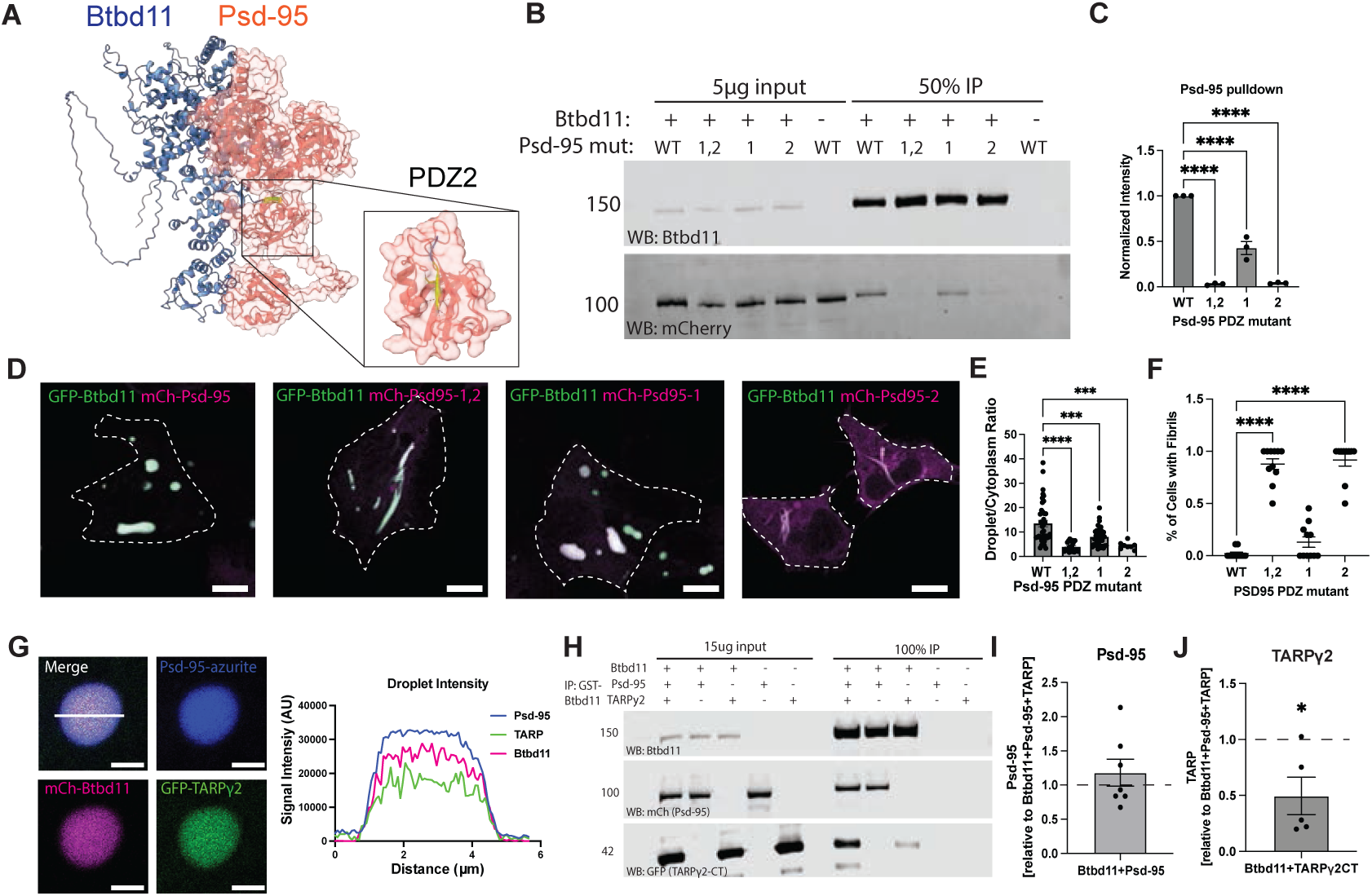
Fine-mapping the Btbd11-PSD-95 interaction and protein complex formation. A) AlphaFold3 prediction of Btbd11 and Psd-95 interaction. B) Representative western blot of GST-Btbd11 and Psd-95-mCherry mutants following GST-Btbd11 pulldown from HEK cells. C) Quantification of Psd-95 mutants show decreased pulldown compared to full-length, one-way ANOVA with Tukey’s multiple comparison < 0.0001, three experimental replicates. D) Visualization of HEK cells transfected with GFP-Btbd11 and Psd-95-mCherry mutants. Scale bar = 10 µm. E) Quantification of Psd-95 colocalized with Btbd11 compared to cytosolic signal ratios show decreased Psd-95 mutant recruitment to Btbd11-containing droplets, one-way ANOVA comparison < 0.0001, three experimental replicates. F) Quantification of percentage of transfected cells containing fibrils, Kruskal-Wallis p < 0.0001, three experimental replicates. G) Cross section of a droplet containing Psd-95-azurite, mCherry-Btbd11, and GFP-TARPγ2-CT shows homogenous colocalization. Scale bar = 2 µm. H) HEK cells were co-transfected with GST-Btbd11, mCherry-Psd-95, and GFP-TARPγ2-CT. Following Btbd11 pulldown, Psd-95 and TARPy2CT were detected. I) Quantification of Psd-95 pulldown band intensity when expressed with Btbd11 alone was normalized to Psd-95 pulldown band intensity of all three proteins expressed together shows no difference. One sample t-test p = 0.8731, 6 replicates. Error bar represents SEM. J) Quantification of TARP pulldown band intensity when expressed with Btbd11 alone was normalized to TARP pulldown band intensity of all three proteins expressed together. One sample t-test, p = 0.0391, 5 biological replicates. Error bar represents SEM.

We further validated our findings using confocal imaging. We co-expressed GFP-Btbd11 with full-length Psd-95-mCherry or PDZ mutants (1, 2, or both) in HEK cells (Figure 1D). Full-length Btbd11 and Psd-95 form homogenous droplet-like condensates when able to interact, in contrast to the fibril-like structure Btbd11 exhibits when expressed alone or unable to interact with other proteins ^32^. When expressed alone, Psd-95 exhibits a cell-fill expression pattern, in comparison to an enrichment in droplet signal intensity when interacting with Btbd11. As expected, full-length Psd-95 effectively forms droplets when interacting with Btbd11. However, both the PDZ2 and combined PDZ1/2 mutants most markedly reduced droplet formation (normalized to the cell cytoplasm; Figure 1E). The PDZ1 mutation alone reduced droplet formation to a lesser extent. In contrast, the proportion of cells containing fibrils rather than droplets, a proxy for disrupted Btbd11-PSD-95 binding, was significantly increased in the PDZ2 and combined PDZ1/2 mutants but not in the PDZ1 mutant (Figure 1F).

Psd-95 regulates AMPA receptor localization at the synapse through interactions with TARPs ^22,30^, of which TARPγ2 is the most abundant isoform in INs ^31^. To determine whether Btbd11 interacts with the Psd-95-TARPγ2 complex, we co-expressed mCherry-Btbd11, Psd-95-azurite, and GFP-TARPγ2 C-terminal fragment (TARPγ2-CT) in HEK cells, which resulted in homogenous, colocalized droplets (Figure 1G). To ensure the droplets reflected a biochemical interaction, we co-expressed GST-Btbd11, Psd-95-mCherry, and GFP-TARPγ2-CT together in HEK cells before lysing and performing GST pulldowns followed by western blotting. When GST-Btbd11, Psd-95-mCherry, and GFP-TARPγ2-CT are expressed together, we found that Btbd11 was able to pull down both Psd-95 and TARPγ2 (Figure 1H). The presence or absence of TARPγ2 did not affect the amount of Psd-95 that was in complex with Btbd11 (Figure 1I). Surprisingly, Btbd11 alone was able to pull down TARPγ2 (Figure 1H), though this occurred to a lesser extent in the absence of Psd-95 (Figure 1J). Together, this data shows Btbd11 and Psd-95 interact via Btbd11’s PDZ-binding motif to PDZ2 of Psd-95, and that TARPγ2 is able to form a protein complex alongside Btbd11 and Psd-95.

### Btbd11 & Psd-95 exist in a phase separated complex with TARP

Given our finding that Btbd11 and TARP form a protein complex with Psd-95, yet both bind via PDZ domains, we next tested whether increasing Btbd11 concentration competes with TARPγ2 for Psd-95 binding. This is important as Psd-95’s function as a synaptic scaffold relies on its ability to bind multiple protein complexes to regulate PSD composition and synaptic plasticity^25,36^. A well-characterized example of this in ENs is SynGAP, a negative regulator of TARP-AMPAR binding to Psd-95, which occupies PDZ “slots” until activity-dependent dispersion from the PSD allows increased binding of the TARP-AMPAR complex ^37,38^. To explore whether similar interactions underlie the Btbd11-Psd-95-TARPγ2 complex, we tested whether Btbd11 competes with TARPγ2 for Psd-95 binding in a SynGAP-like manner. We co-expressed consistent equimolar amounts of Psd-95-azurite and GFP-TARPγ2-CT with increasing amounts of mCherry-Btbd11 in HEK cells before measuring the signal of each component in droplets compared to the cytoplasmic signal (Figure 2A, individual channels displayed in Supplemental Figure 2A). We noted that these droplets were reminiscent of a liquid-liquid phase separated state, as seen previously with Btbd11 and Psd-95 ^32^ and Psd-95 with other glutamatergic PSD proteins ^13,39^. We found the amount of GFP-TARPγ2-CT is not impacted by the amount of Btbd11, a notable difference from a similar experiment performed by the Huganir lab in which SynGAP outcompetes TARP binding to Psd-95 in a dose-dependent manner ^38^. We found that higher Btbd11 signal led to greater Psd-95 signal enrichment in droplets and increased overall droplet size (Figure 2B). We confirmed our results by co-expressing Psd-95-GST and GFP-TARPγ2-CT with increasing concentration of mCherry-Btbd11 and performing an Psd-95-GST IP (Figure 2C). TARPγ2-CT binding to Psd-95 was unchanged regardless of increasing Btbd11 concentrations, suggesting Btbd11 does not prevent TARPγ2 from binding to Psd-95 when all three are present a protein complex. In fact, inclusion of Btbd11 compared to Psd-95 and TARPγ2-CT alone increases the average size of observed droplets (Figure 2D).

**Figure 2.**
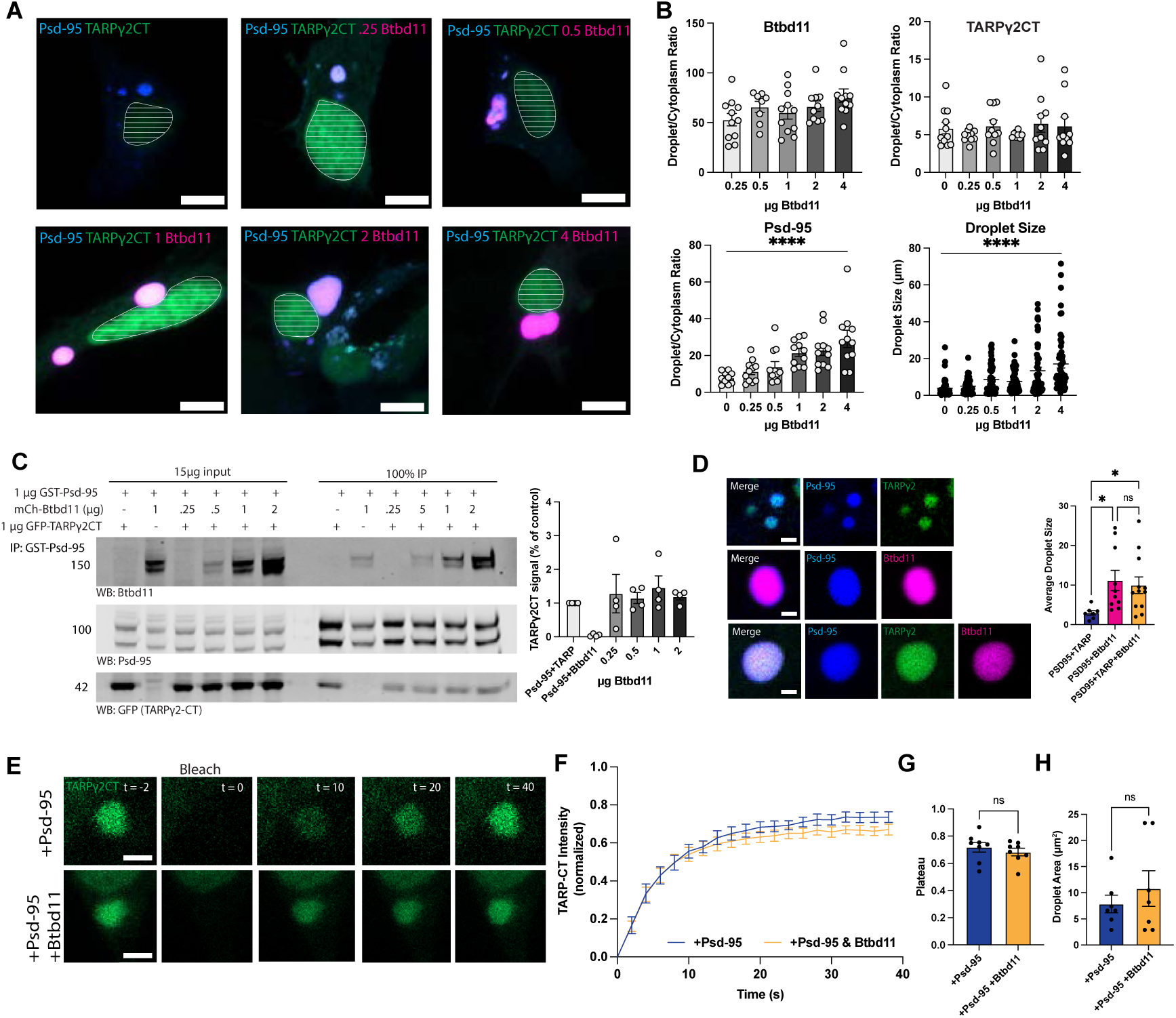
Btbd11, PSD-95, and TARPγ2 form a phase-separated protein complex. A) HEK cells were co-transfected with azurite-Psd-95, GFP-TARPγ2-CT, and increasing concentration of mCherry-Btbd11. Shapes with horizontal lines represent GFP-TARPγ2-CT localized to the nucleus and were not included in analysis. Scale bar = 10 µm. B) Ratio of intensity in droplet to cytoplasm was measured for each component, showing no significant effect of Btbd11 concentration on Btbd11 or TARPγ2 recruitment (one-way ANOVA p = 0.0878 and 0.68, respectively). PSD-95 localization to droplets over cytoplasm increased with Btbd11 (one-way ANOVA p<0.0001), and average condensate size increased with increased Btbd11 concentration (Kruskal-Wallis test p = 0.0009). N = 11 cells per condition from 3 biological replicates. C) HEK cells were co-transfected with GST-Psd-95, GFP-TARPγ2-CT, and increasing concentration of mCherry-Btbd11. Following Psd-95 pulldown, Btbd11 and TARPy2CT were detected. Quantification of TARPγ2-CT pulldown shows no significant effect of Btbd11 concentration, 4 biological replicates. D) Droplets containing Btbd11 are significantly larger than those without Btbd11 (Welch’s one-way ANOVA p = 0.0027 with Dunnett’s multiple comparisons test, 3 biological replicates) Scale bar = 1µm. E) Representative confocal images show photobleaching of TARPγ2-CT in droplets containing PSD-95 with and without Btbd11 over time. Scale bar = 2µm. F) One-phase association best-fit lines of TARPγ2-CT recovery in droplets containing PSD-95 and Btbd11 shows decreased recovery over time compared to TARPγ2-CT and PSD-95 alone. G) Comparison of TARPγ2-CT in droplets containing Psd-95 and Btbd11 or Psd95 alone show no difference in plateau. Unpaired t-test p = 0.442, n = 8 droplets from 3 batches of cells. H) Comparison of droplet size used for analysis in E-G. Welch’s t-test p = 0.4579, n = 8 droplets from 3 batches of cells. Error bars represent SEM.

To show dynamic recovery over time, a hallmark property of phase separation, we performed fluorescent recovery after photobleaching (FRAP) of TARPγ2-CT in droplets with Psd-95 and Btbd11 or with Psd-95 alone, showing dynamic recovery in each condition (Figure 2E). Recovery kinetics of TARPγ2-CT after photobleaching were similar whether Btbd11 was included or not, suggesting similar TARPγ2-CT mobility in each condition (Figure 2F-G). Droplets of similar size were chosen for bleaching to avoid potential confounding effects of size on recovery speed (Figure 2H).

We next used FuzDrop ^40^, a sequence-based algorithm that predicts the propensity of a given protein to undergo phase separation, to determine the probability of Btbd11 as a driver of LLPS. Indeed, Btbd11 had a probability of LLPS (pLLPS) of 0.9712 (Supplemental Figure 2B), and as proteins with pLLPS above 0.60 are likely promoters of LLPS, we expect Btbd11 is a driver of LLPS with Psd-95 and TARPγ2-CT. To demonstrate droplets containing all three proteins display another property of LLPS, we showed puncta coalescence over time with live-cell confocal imaging (Supplemental Figure 2C). We then predicted the propensity of Btbd11 lacking the N-terminal domain to phase separate using FuzDrop, finding a pLLPS of 0.3305 (Supplemental Figure 2D), in alignment with a previously documented observed inability to phase separate ^32^. We next demonstrated co-expression of Psd-95 and TARPγ2-CT with Btbd11ΔN form aggregates compared to droplets (Supplemental Figure 2E), which show dramatically reduced TARPγ2-CT recovery when photobleached (Supplemental Figure 2F-H), consistent with protein binding and unbinding, rather than more dynamic exchange expected with LLPS.

Collectively, these findings establish Btbd11 as a component of the Psd-95/TARP complex capable of driving phase separation through discrete protein domain interactions.

### Btbd11 synapse localization

We previously found that Btbd11 is primarily localized to glutamatergic synapses in INs ^32^. To confirm these findings, we performed immunostaining on primary cultured rat hippocampal neurons with GAD67 (to label INs) and VGlut1 (to label glutamatergic presynaptic terminals) along with Btbd11 and Psd-95 (Figure 3A). We observed that Btbd11 is found at nearly 80% of VGlut1-positive synapses that contain Psd-95 puncta within GAD67-positive interneurons (Figure 3B). Further, synaptic Btbd11 and Psd-95 intensity within IN VGlut1-positive synapses exhibited a positive correlation (Figure 3C).

**Figure 3.**
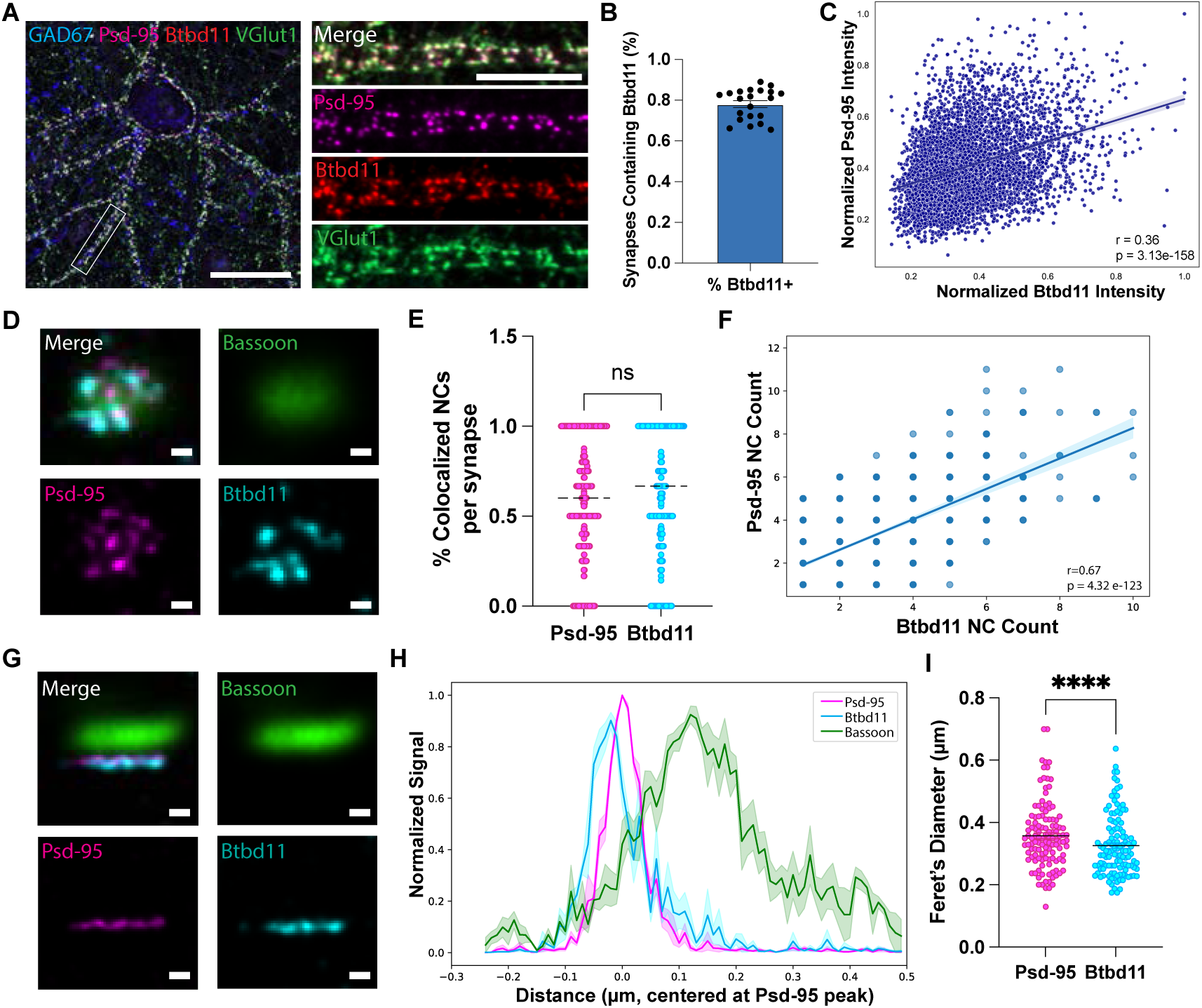
Btbd11 synaptic localization and nanostructure. A) Primary rat hippocampal neurons stained for GAD67, Btbd11, PSD-95, and VGlut1, scale bars = 30 and 10µm, respectively. B) Percentage of VGlut1 and Psd-95 synapses containing Btbd11, mean = 0.7813, n = 21 neurons from 3 independent batches of cells. Error bars represent SEM. C) Pearson correlation of synaptic Btbd11 and PSD-95 intensity. r = 0.37, p = 3.13 e-158, n = 5073 synapses from 3 batches of neurons. Shading represents SEM. D) Representative image of en face synapse, showing confocal Bassoon with STED Psd-95 and Btbd11. Scale bar = 0.01µm. E) Comparison of percentage of colocalized Psd-95 and Btbd11 NCs per synapse. Mann-Whitney test p = 0.2037, Psd-95 n = 953 synapses, Btbd11 n = 991 synapses from 3 batches of neurons. F) Correlation between Psd-95 and Btbd11 nanocluster number per synapse. Pearson r = 0.67, p = 4.32 e-12, n = 948 synapses from 3 batches of neurons. Shading represents SEM. G) Representative image of side synapse, showing confocal Bassoon with STED Psd-95 and Btbd11. Scale bar = 0.01µm. H) Normalized average maximum intensity across synapse line scan, n = 93 synapses from 3 batches of neurons. Error bars represent SEM. I) Feret’s diameter of Psd-95 and Btbd11, Mann-Whitney test p = 0.0056, n = 93 synapses from 3 batches of neurons. Error bars represent SEM.

Synaptic nanostructure is generally conserved between ENs and INs, with Psd-95 in interneurons displaying distinct clustering and pre-synaptic alignment consistent with established nanocolumn features ^6^. We thus used stimulated emission depletion (STED) microscopy to examine the precise synaptic localization of Btbd11 relative to Psd-95 at a super resolution level in primary cultured rat hippocampal neurons (Figure 3D; raw STED channels shown in Supplemental 3A). We found Btbd11 formed distinct nanoclusters (NCs), rather than being uniformly distributed across the PSD. To examine differences in Psd-95 and Btbd11 NC properties, we analyzed images with a custom Fiji detection pipeline in which NCs in *en face* synapses, where NCs can be identified with highest fidelity^41^, were each defined by unsharp masking^42^, as described in *Methods* (representative synapse and NC detections shown in Supplemental Figure 3B). We validated that Psd-95 NC count per *en face* synapse positively correlates with synapse size as expected in mature neuron PSDs ^6,21,43^, and discovered Btbd11 exhibits a similar positive correlation (Supplemental Figure 3C). Noting distinct Psd-95 and Btbd11 NC patterns, we analyzed the colocalization of Btbd11 and Psd-95 NCs. 60% of Psd-95 and Btbd11 NCs were colocalized with a NC of the other protein (Figure 3E). Colocalized Psd-95 and Btbd11 NCs showed a broad distribution of intersection over union values, indicating that even when co-localized, they occupy distinct nanoscale distributions within the synapse (Supplemental Figure 3D). Consistent with this, while on average more than half the NCs at a given synapse were colocalized, the variance was broad, and many synapses exhibited either 100% colocalization or no NC colocalization (examples shown in Supplemental Figure 3E and F, respectively). We next examined the relationship between Psd-95 and Btbd11 NCs, finding that the number and size of Btbd11 NCs positively correlates with Psd-95 per synapse (Figure 3F, Supplemental Figure 3G), and overall, the median Btbd11 NC size was significantly larger than Psd-95 NCs by 7.41% (Supplemental Figure 3H).

At synapses viewed from the side, we found Btbd11 signal was consistently offset deeper into the PSD relative to Psd-95 and presynaptic marker Bassoon (Figure 3G, raw STED channels in Supp. 3I). To quantify this, we measured signal intensity for each protein along a line scan perpendicular to the PSD. Btbd11 and Psd-95 showed distinct average normalized maximum peak signals along a line scan (Figure 3). Consistent with previously published studies^44,45^, Psd-95 and Bassoon displayed a transsynaptic distance on average of 120 nanometers (Supplemental Figure 3J), and we found Btbd11 mean peak intensity was measured approximately 25 nm below Psd-95. Further, we found that Btbd11 had a significantly shorter average width at the PSD than Psd-95, as measured by Feret’s diameter (Figure 3I), which may suggest a more centralized synaptic localization. It is important to note that the Btbd11 antibody used binds at the N-terminus, which is expected to be found farther away from the Psd-95 binding site based on protein interaction models (Figure 1A). Thus, the STED signal likely reflects the localization of Btbd11’s N-terminal region, which is responsible for LLPS, though the exact conformation of Btbd11’s N-terminus relative to its PDZ-binding C-terminus remains unknown.

Together, these findings show that Btbd11 and PSD-95 occupy overlapping but distinct nanoscale distributions, a pattern consistent with their ability to directly interact. Given that Btbd11 is present at nearly all glutamatergic synapses, these data position it as a putative regulator of excitatory synaptic function.

### Btbd11 expression impacts post-synaptic protein stability in a phase separation-dependent manner

Having shown the precise localization of Btbd11 adjacent to Psd-95 at the PSD, we speculated it was well-positioned to regulate the stability of synaptic proteins. Btbd11’s N-terminus is essential for LLPS in HEK cells, so to address this question, we tested whether deletion of the N-terminus affects Btbd11’s ability to regulate protein stability in INs. First, we characterized the synaptic stability of full-length Btbd11 compared to the phase separation-null mutant, Btbd11ΔN. We expressed mCherry-Btbd11 or mCherry-Btbd11ΔN alongside genetically encoded Psd-95 intrabody Psd-95.FingR-GFP ^46^ as a synaptic marker in putative INs identified via mDlx-azurite and performed FRAP on Btbd11 (Figure 4A). We found a dramatic reduction in the recovery kinetics of Btbd11ΔN compared to full-length Btbd11 (Figure 4B-C), indicating a very large immobile fraction, consistent with the hypothesis that an inability to phase separate reduced protein mobility in the crowded PSD.

**Figure 4.**
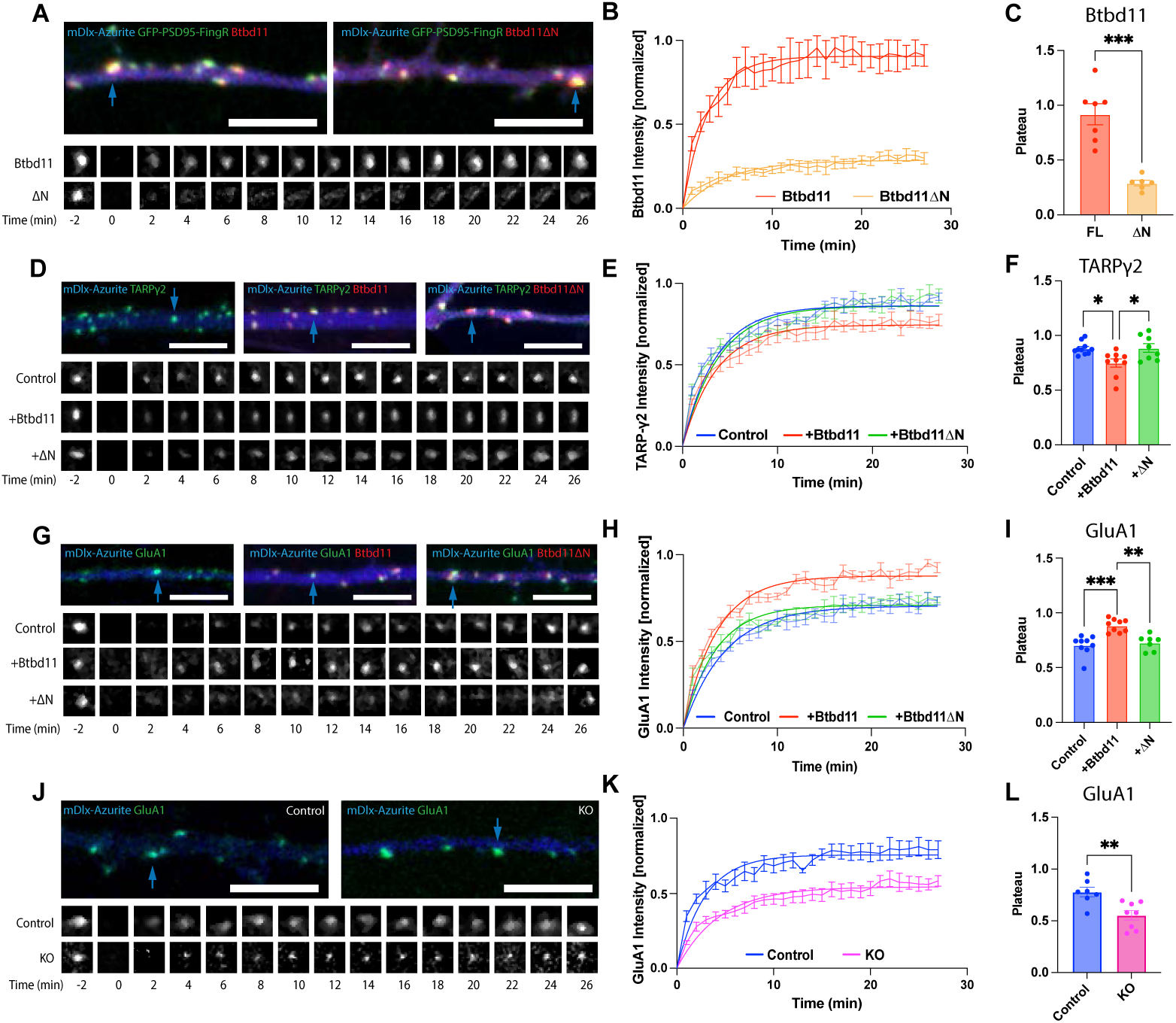
Btbd11 expression and phase separation impact glutamatergic PSD protein stability. A) Representative primary cultured rat hippocampal neurons transfected with mDlx-azurite and GFP-Psd-95-FingR and mCherry-Btbd11 or mCherry-Btbd11ΔN. Lower panels show FRAP of puncta indicated by blue arrow in each condition. B) Recovery of Btbd11 with one-phase association best fit line. C) Quantification of estimated plateau value from best-fit line, unpaired t-test with Welch’s correction p = 0.0004, n = 6-7 cells from 6 independent cultures. Error bars display SEM. D) Representative primary cultured rat hippocampal neurons transfected with mDlx-azurite and GFP-TARPγ2 alone, with mCherry-Btbd11, or mCherry-Btbd11ΔN. Lower panels show FRAP of puncta indicated by blue arrow in each condition. E) Recovery of TARPγ2 with one-phase association best fit line. F) Quantification of estimated plateau value from best-fit line, one-way ANOVA F = 5.776, p = 0.009, n = 8-10 cells from 5 independent cultures. Error bars display SEM. G) Representative primary cultured rat hippocampal neurons transfected with mDlx-SEP-GluA1 and mDlx-Azurite alone, with mCherry-Btbd11, or mCherry-Btbd11ΔN. Lower panels show FRAP of puncta indicated by blue arrow in each condition. H) Recovery of GluA1 with one-phase association best fit line. I) Quantification of estimated plateau value from best-fit line, one-way ANOVA F = 15.01, p < 0.0001, n = 7-9 cells from 4 independent cultures. Error bars display SEM. J) Representative primary cultured rat hippocampal neurons transfected with mDlx-SEP-GluA1 and mDlx-Azurite in control or CRISPR-mediated Btbd11 knockout. Lower panels show FRAP of puncta indicated by blue arrow in each condition. K) Recovery of GluA1 with one-phase association best fit line. L) Quantification of estimated plateau value from best-fit line, unpaired t-test p = 0.0043, n = 7-8 cells from 4 independent cultures. Error bars display SEM. All scale bars = 5µm.

We previously showed that full-length Btbd11 was able to stabilize Psd-95 at IN synapses ^32^. We next examined the interaction between Btbd11 and SAP102 to see if the previously demonstrated stabilization and interaction with Psd-95 was specific or generalized to other MAGUKs. SAP102 is a highly mobile Psd-95 homolog, with similar yet distinct synaptic properties and localization, and had been previously identified as a putative Btbd11 interactor ^32,47,48^. Similar to Psd-95, Btbd11’s PBM was predicted by AlphaFold3 to bind to SAP102’s PDZ2 (Supplemental Figure 4A). We confirmed the interaction between Btbd11 and SAP102 using co-IP in HEK cells (Supplemental Figure 4B). Further, confocal imaging of co-expressed Btbd11 and SAP102 in HEK cells forms colocalized putative condensates similar to those seen with Psd-95 (Supplemental Figure 4C). We demonstrated that binding was absent when Btbd11’s PBM is deleted (Supplementary Figure 4D-E). Btbd11’s N-terminus, in contrast to the PBM, is not necessary for SAP102 binding (Supplementary Figure F-G). To investigate whether Btbd11 overexpression stabilizes SAP102, we expressed mCherry-SAP102 with mDlx-azurite alone, with GFP-Btbd11, or GFP-Btbd11ΔN in rat hippocampal neurons and performed FRAP on SAP102 (Supplemental Figure 4H). Consistent with Btbd11’s stabilization effect previously seen on Psd-95, we found a similar slowed recovery of SAP102 with Btbd11 overexpression compared to control (Supplemental Figure 4K). Interestingly, we saw no stabilization effect with ΔN overexpression, with recovery kinetics indistinguishable from the Control condition (Supplemental Figure 4I). There was a significantly lower maximum recovery, or plateau, of SAP102 following Btbd11 overexpression compared to control and ΔN (Supplemental Figure 4J).

Next, we tested the effect of Btbd11 and ΔN overexpression on TARPγ2 stability. We wondered whether we would see a similar stabilization effect, given the inclusion of TARPγ2 in the Psd-95 complex and reliance on Psd-95 for synaptic clustering. We expressed GFP-TARPγ2 with mDlx-azurite alone, with mCherry-Btbd11, or mCherry-Btbd11ΔN and performed FRAP on TARPγ2 (Figure 4D). We observed a stabilizing effect of Btbd11 overexpression on TARPγ2 recovery, but no effect with the ΔN mutant (Figure 4E-F), again highlighting the requirement of Btbd11 to be able to undergo LLPS in order to impact the mobility of other synaptic proteins.

We then directly examined the impact of Btbd11 overexpression on AMPAR stability in INs using fluorescently tagged GluA1. To specifically look at surface AMPAR in interneurons, we co-expressed mDlx-SEP-GluA1 alone with mDlx-azurite, with mCherry-Btbd11, or with mCherry-Btbd11ΔN and performed FRAP on GluA1 (Figure 4G). We found a surprising increase in GluA1 recovery kinetics with Btbd11 overexpression compared to both control and ΔN, though consistently, no difference between control and ΔN (Figure 4H-I). We confirmed that the effect of Btbd11 overexpression on GluA1 recovery in synapses was indirect, as an IP revealed Btbd11 does not bind to GluA1 directly (Supplemental Figure 4K).

Next we sought to test if Btbd11’s effect on GluA1 mobility was bi-directional, using CRISPR/Cas9 mediated Btbd11 KO in primary rat hippocampal neurons (Figure 4J). We first confirmed Btbd11 knockdown efficiency using co-expression of GFP-Btbd11 with or without the CRISPR construct in HEK cells (Supplemental Figure 4L). Reassuringly, Btbd11 knockout (KO) led to decreased SEP-GluA1 recovery compared to control (Figure 4K-L), showing a bi-directional effect of Btbd11 expression on GluA1 stability. Together, these data show that Btbd11 affects the stability of key synaptic proteins responsible for glutamatergic synaptic transmission in a phase separation dependent manner

### Btbd11 KO leads to a reduction in synaptic protein expression *in vitro*

Given Btbd11’s role in PSD protein stability, we next asked whether Btbd11 KO impacts the abundance of proteins at IN glutamatergic synapses. For the following experiments, we cultured primary cultured Btbd11^FL/FL^ mouse neurons, generating control or KO neurons via transduction with AAV-mCherry or AAV-mCherry-Cre. At DIV14-15, cells were PFA-fixed and stained for GAD67 (to identify INs), VGlut1 (presynaptic marker), and either Psd-95, TARPγ2, or GluA1 prior to confocal imaging. Images were processed and analyzed using SynAPSeg, an open-source deep learning-based synapse analysis pipeline ^49^ (Supplemental Fig 5A-C). We first examined how Psd-95 abundance was altered in Btbd11 KO neurons (Figure 5A). We quantified Psd-95 that colocalized with VGlut1 in GAD67-positive neurons, finding significantly decreased synaptic Psd-95 puncta size and density in KO cells compared to control (Figure 5B-C). Next, we investigated the impact of Btbd11 KO on TARPγ2 abundance (Figure 5D). Similar to Psd-95, we quantified changes in TARPγ2 puncta colocalized with VGlut1 in GAD67-positive neurons. We found significantly decreased synaptic TARPγ2 intensity and density in the KO condition compared to control (Figure 5E-F). Finally, we investigated the impact of Btbd11 KO on AMPARs by quantifying GluA1 abundance in Btbd11 KO neurons compared to control (Figure 5G). We quantified changes in GluA1 puncta colocalized with VGlut1 in GAD67-positive neurons, finding significantly decreased synaptic GluA1 puncta intensity and density in the KO condition (Figure 5H-I). Collectively, these data show that Btbd11 KO has a striking impact on synaptic expression and density of PSD proteins essential for glutamatergic signaling.

**Figure 5.**
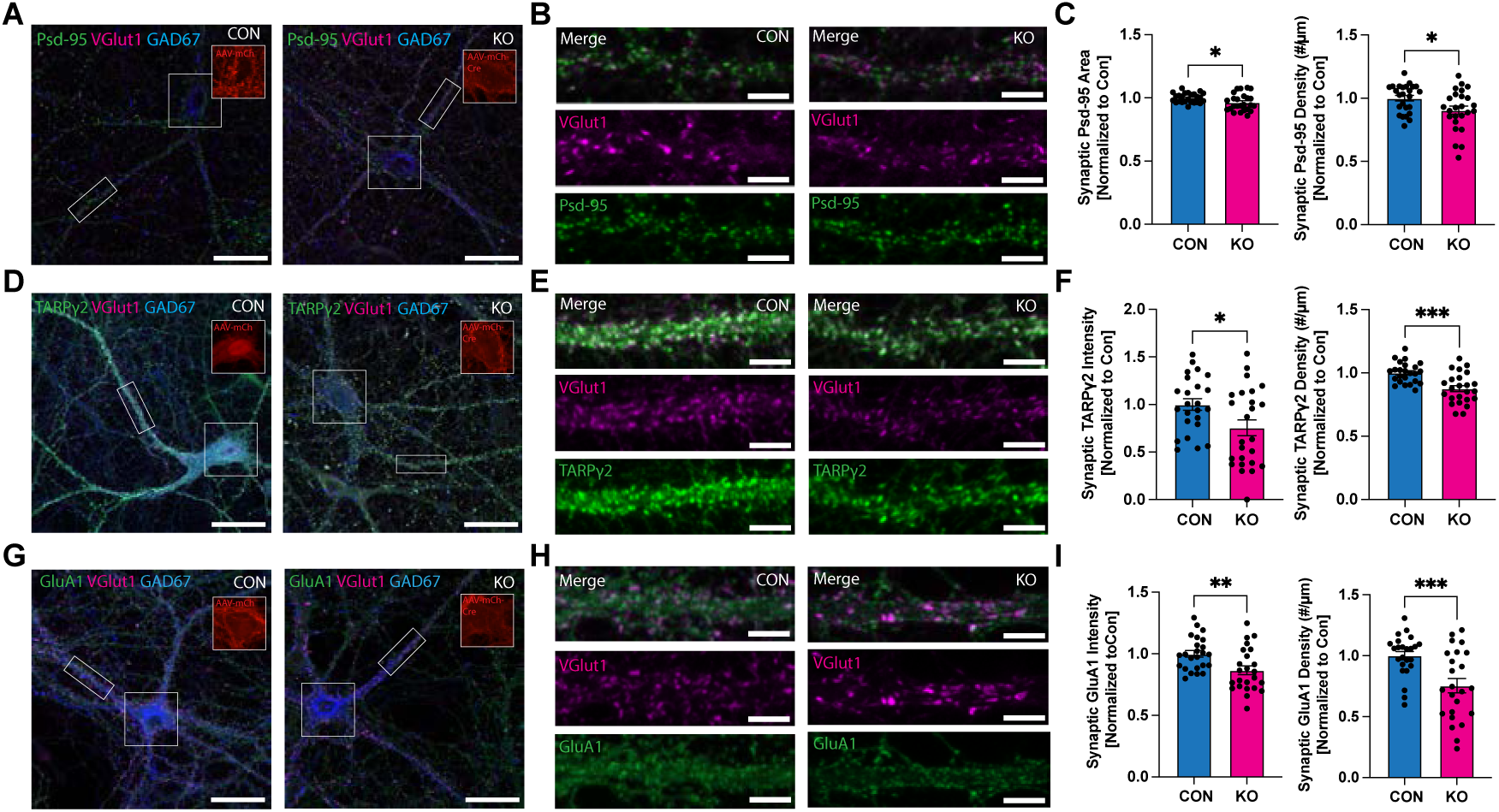
Btbd11 knockout impacts synaptic protein expression in vitro. A) Representative confocal Control and Btbd11 knockout neurons stained for Psd-95, VGlut1, and GAD67. Scale bars represent 30µm. B) Representative dendrite section showing synaptic PSD-95 colocalized with VGlut1. Scale bars represent 5µm. C) Btbd11 knockout neurons show significantly decreased puncta area (Welch’s t-test, p = 0.0242) and density (unpaired t-test, p = 0.0202) compared to control. n = 25 neurons from 4 independent cultures. D) Representative Control and Btbd11 knockout neurons stained for TARPγ2, VGlut1, and GAD67. Scale bars represent 30µm. E) Representative dendrite section showing synaptic TARPγ2 colocalized with VGlut1. Scale bars represent 5µm. F) Btbd11 knockout neurons show significantly decreased puncta area (unpaired t-test, p = 0.021) and density (unpaired t-test, p = 0.0001) compared to control. n = 24 neurons from 4 independent cultures. G) Representative Control and Btbd11 knockout neurons stained for GluA1, VGlut1, and GAD67. Scale bars represent 30µm. H) Representative dendrite section showing synaptic GluA1 colocalized with VGlut1. Scale bars represent 5µm. I) Btbd11 knockout neurons show significantly decreased puncta intensity (unpaired t-test, p = 0.0049) and density (Welch’s t-test, p = 0.0009) compared to control. n = 24 neurons from 4 independent cultures. All error bars represent SEM.

### Btbd11 KO leads to reduced spontaneous calcium transients

We next investigated whether the decreased expression of postsynaptic glutamatergic PSD proteins in Btbd11 KO neurons would manifest as a functional change in signaling. Spontaneous calcium transients provide a similar readout compared to electrophysiological EPSC recordings ^50^, so we generated Btbd11 control and KO neurons through viral mCherry or mCherry-Cre delivery in primary cultured Btbd11^FL/FL^ mouse neurons. This was followed by subsequent transduction with AAV-mDlx-GCamp8f, to enable the specific visualization of calcium transients in INs. We used pharmacological blockade of action potentials with tetrodotoxin and inhibition of NMDARs using AP5 to isolate spontaneous AMPAR mediated calcium transients in INs as a proxy for miniature excitatory postsynaptic currents (Figure 6A-B). We visualized calcium transients (ΔF/F) within dendrites in control (Figure 6C) and Btbd11 KO cells (Figure 6D). By aligning calcium events across genotype (Figure 6E), we found that Btbd11 KO in INs have reduced event amplitudes (Figure 6E-F), as well as a dramatically reduced event frequency (Figure 6H), consistent with prior electrophysiological recordings in the visual cortex of IN-specific Btbd11 KO mice ^32^. We noted a significant difference between control and knockout event frequency prior to NMDAR blockade (Supplementary Figure B), though there was no change in event frequency within either genotype condition after NMDAR blockade, showing no major contribution of NMDARs to spontaneous events (Supplementary Figure C-D). Finally, to confirm the recorded events were from AMPARs, we pharmacologically blocked AMPAR with DNQX application and observed a near complete signa drop-off (Supplementary Figure B). Together, these data show a functional postsynaptic deficit in calcium signaling resulting from loss of Btbd11 in glutamatergic synapses.

**Figure 6.**
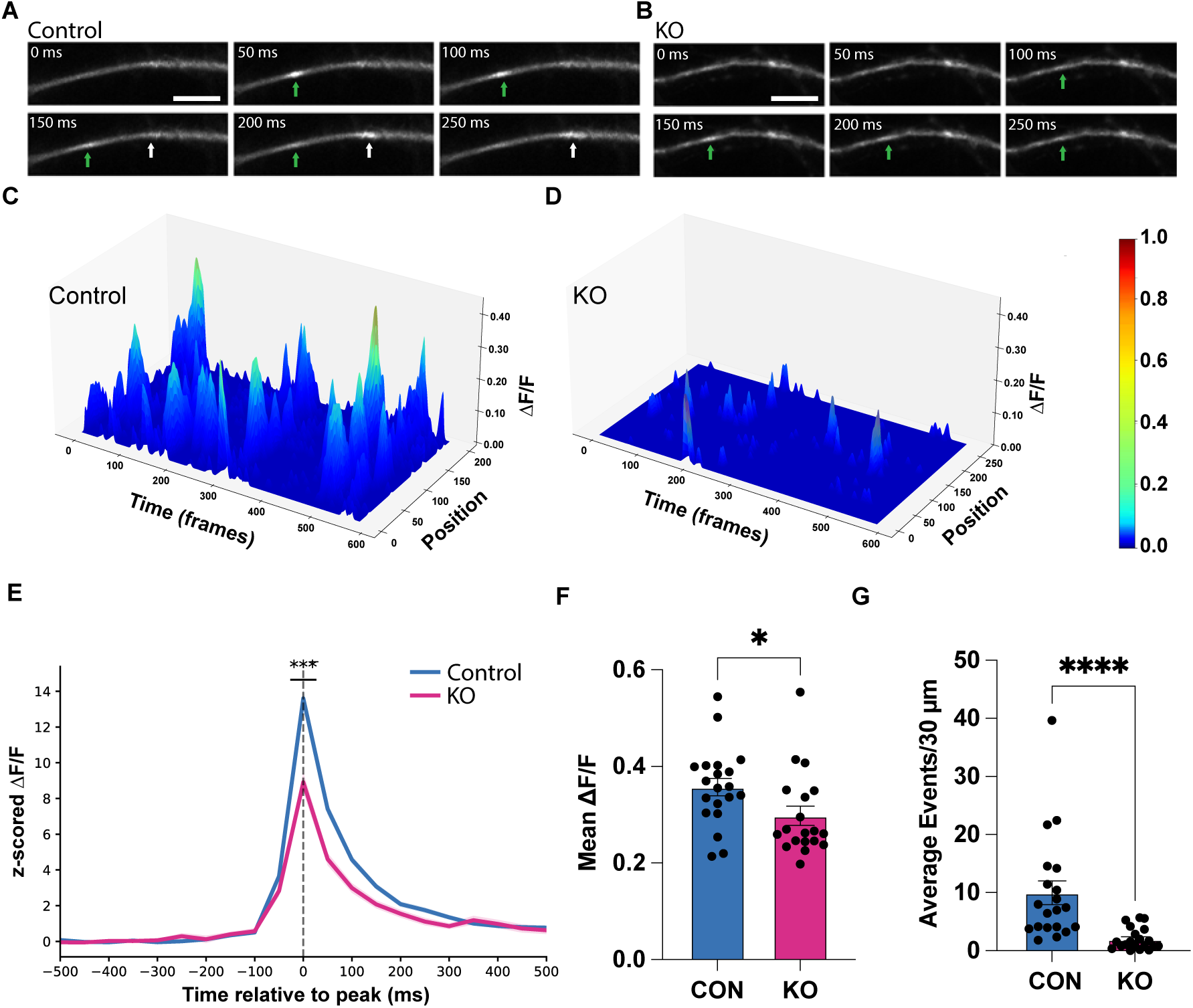
Spontaneous calcium signaling is diminished in Btbd11 knockout neurons. A) Representative timelapse imaging of spontaneous AMPAR-mediated calcium transients in putative interneurons from cultured mouse hippocampal control neurons. Cultures treated with TTX to block action potentials and AP5 to block NMDARs. Arrows indicate calcium event. Scale bar = 50µm. B) Representative timelapse imaging of spontaneous calcium transients in putative interneurons from cultured mouse hippocampal Btbd11 knockout neurons. Arrows indicate calcium event. Scale bar = 50µm. C) Representative calcium events from one control dendrite over time. D) Representative calcium events from one Btbd11 knockout dendrite over time. E) Averaged z-scored ΔF/F signal for all control and knockout events. Welch’s t-test p < 0.0001, Control n = 514 events, KO n = 103 events. F) Comparison of control and Btbd11 KO mean ΔF/F averaged per neuron. Mann-Whitney test p = 0.0262, n = 20 neurons per condition from 3 batches of cells. G) Comparison between average number of calcium events averaged per 30µm between control and KO. Mann-Whitney test p < 0.0001, n = 20 neurons per condition from 3 batches of cells.

### Btbd11 KO leads to reduced Psd-95 nanoscale clustering in interneurons

Finally, to determine whether Btbd11-mediated loss in glutamatergic signaling was related to a loss of nanostructure, we employed STED microscopy to investigate whether Psd-95 NCs are disrupted in Btbd11 KO neurons. Using primary cultured hippocampal Btbd11^FL/FL^ neurons transduced with either AAV-GFP or AAV-GFP-Cre, we imaged 1-channel Psd-95 STED alongside confocal-resolution Bassoon as a designated presynaptic marker. We first replicated our previous results (Figure 5A-C) and observed, again, that Btbd11 KO reduces synaptic Psd-95 area as well as puncta count (Supplementary Figure 7A-B).

Next, we quantified the properties of Psd-95 NCs per *en face* synapse (Figure 7A-B). We found a similar positive correlation between synapse size and NC count per synapse regardless of Btbd11 expression (Supplemental Figure 7C). We then compared the average number of Psd-95 NCs in control compared to Btbd11 KO synapses, finding a significant decrease in the Btbd11 KO condition when NCs per synapse were averaged per synapse (Figure 7C) and with cell averaging (Figure 7D). Similarly, Psd-95 NC area was reduced in Btbd11 KO compared to control, both when averaged per NC (Figure 7E) or per cell (Figure 7F).

**Figure 7.**
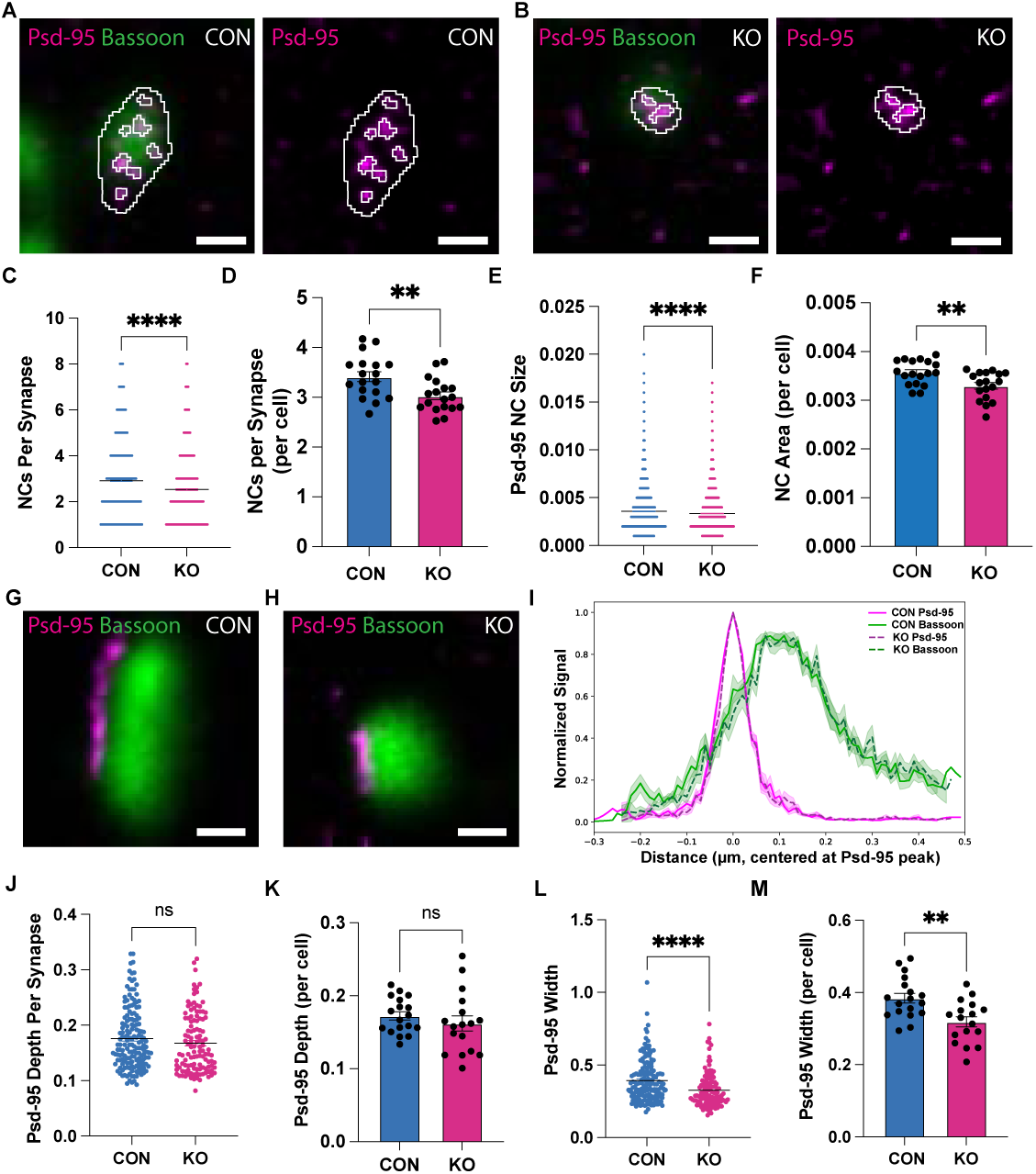
Btbd11 loss disrupts Psd-95 nanostructure. A) Representative Psd-95 *en face* synapse image showing confocal Bassoon with STED Psd-95 and Btbd11, with nanocluster detection outlines from Control neurons. Scale bar = 0.2µm. B) Representative Psd-95 *en face* synapse image showing confocal Bassoon with STED Psd-95 and Btbd11, with nanocluster detection outlines from KO neurons. Scale bar = 0.2µm. C) Comparison between mean number of Psd-95 nanoclusters per synapse averaged by cell, unpaired t-test = 0.0051, n = 18 cells from 3 batches of neurons. Error bars represent SEM. D) Comparison between number of Psd-95 nanoclusters per synapse, Mann-Whitney test p<0.0001, Control n = 2363 synapses, KO n = 1863 synapses. Error bars represent SEM. E) Comparison between mean Psd-95 nanocluster area per synapse averaged by cell, unpaired t-test = 0.0045, n = 18 cells from 3 batches of neurons. Error bars represent SEM. F) Comparison between number of Psd-95 nanoclusters per synapse, Mann-Whitney test p<0.0001, Control n = 2363 synapses, KO n = 1863 synapses. Error bars represent SEM. G) Representative on side synapse Psd-95 STED image with confocal Bassoon and nanocluster detection from Control neurons. Scale bar = 0.2µm. H) Representative on side synapse Psd-95 STED image with confocal Bassoon and nanocluster detection from KO neurons. Scale bar = 0.2µm. I) Representative mean line scan through Psd-95 and Bassoon, with maximum peak value normalized per synapse, Control n = 163 synapses, KO n = 124 synapses. J) Comparison between mean Psd-95 depth averaged by cell, unpaired t-test = 0.3867, n = 17-18 cells from 3 batches of neurons. Error bars represent SEM. K) Comparison of Psd-95 depth per synapse, Mann-Whitney test p = 0.1539, Control n = 162 synapses, KO n = 118 synapses from 3 batches of neurons. Error bars represent SEM. L) Comparison between mean Psd-95 width averaged by cell, unpaired t-test = 0.0021 n = 17-18 cells from 3 batches of neurons. Error bars represent SEM. M) Comparison of Psd-95 width per synapse, Mann-Whitney test p<0.0001, Control n = 163 synapses, KO n = 124 synapses from 3 batches of neurons. Error bars represent SEM.

Side synapses (Figure 7G-H) revealed no change in the distribution of Psd-95 signal between control and KO conditions (Figure 7I), or any change in the depth of the Psd-95 signal Psd-95, or thickness of Psd-95 away from the membrane, either individual synapse or by cell (Figure 7J-K). However, we did see a significant decrease in Psd-95 width in Btbd11 KO compared to control, as measured by Feret’s diameter, both when averaged total synapses (Figure 7L) or by cell (Figure 7M), consistent with the observed reduction in PSD area. Together, these data provide evidence that Btbd11 plays a role in nanoscale Psd-95 configuration, specifically by supporting the ability to maintain both the size and number of Psd-95 NCs.

## Discussion

In this study, we use a combination of biochemistry, live cell imaging, and super resolution imaging to demonstrate that Btbd11 supports glutamatergic signaling in hippocampal INs through specific interactions with essential PSD proteins and the promotion of phase separation at the synapse. We further show that Btbd11 KO leads to a loss of AMPAR mobility, synaptic protein expression, Psd-95 NC number and size, and glutamatergic signaling in inhibitory INs (Figure 8).

**Figure 8.**
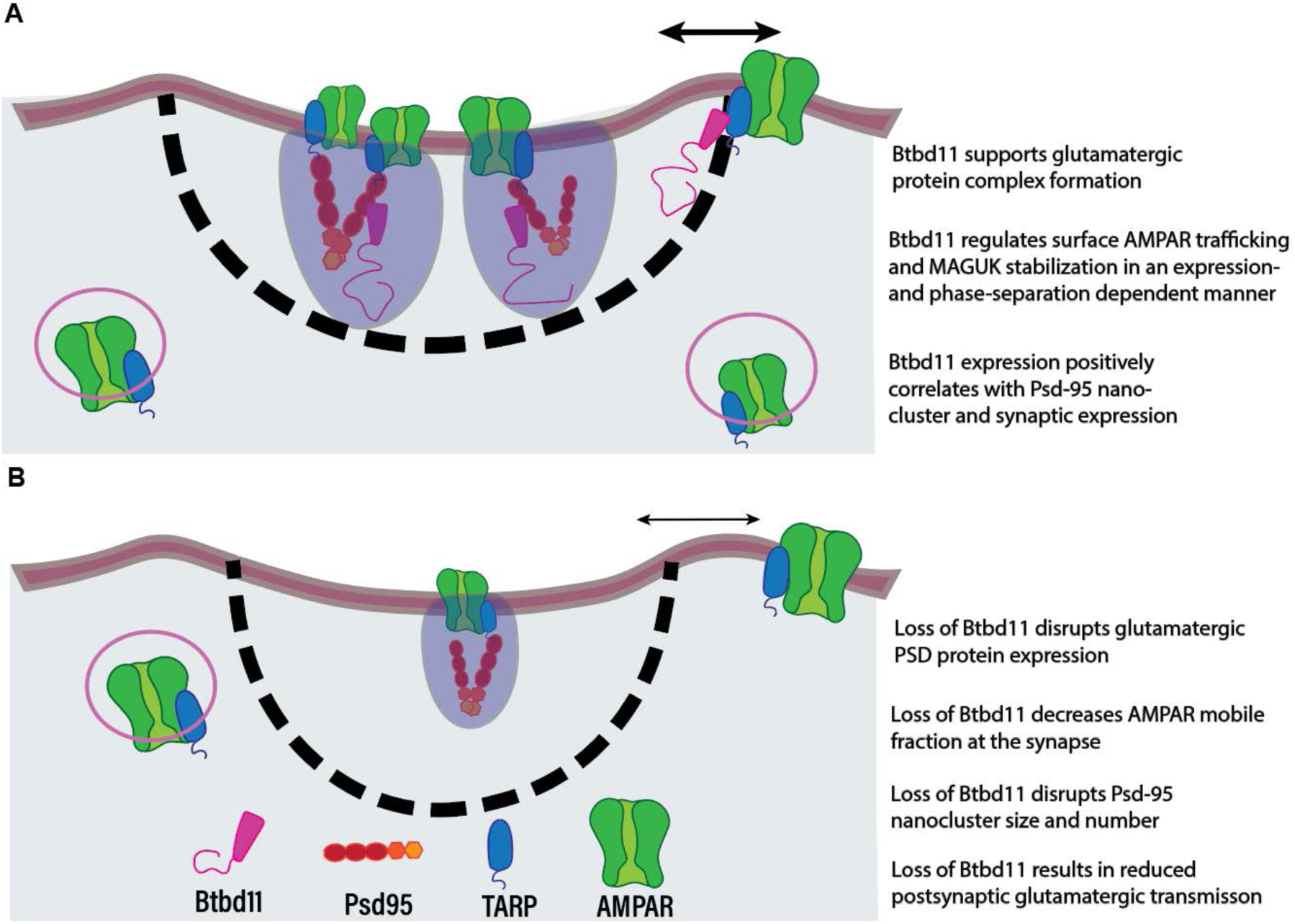
Schematic of Btbd11 mechanism at the synapse. A) Btbd11 supports glutamatergic transmission in INs through regulating PSD protein expression and mobility at the synapse. B) Btbd11 knockout disrupts PSD protein expression and mobility in INs, resulting in impaired glutamatergic transmission.

Given the abundance and importance of Psd-95 as a regulator of synaptic transmission via scaffolding with a variety of receptors and signaling molecules, understanding the interactions it forms with distinct binding partners is crucial. As Btbd11 has previously been detected within the Psd-95 complex ^32,36^ and established as a supporter of glutamatergic transmission, we sought to identify the specific interaction site. We found that Btbd11 preferentially binds to Psd-95 PDZ2, the same domain that TARPγ2 is known to bind to^30^. We chose to focus specifically on protein complex formation of Btbd11 and Psd-95 with TARPγ2, the most abundant TARP isoform in interneurons ^31^, because of its critical role anchoring AMPARs at the synapse and subsequent mediation of glutamatergic transmission. We found that Psd-95, Btbd11, and TARPγ2 all colocalize in putative phase-separated condensates. Curiously, TARPγ2-CT can directly interact with Btbd11, though the interaction was weaker alone compared to when in complex with both Btbd11 and Psd-95. Inclusion of Btbd11 into this complex establishes the molecular basis through which it is positioned to mediate glutamatergic signaling in INs, as discussed below.

### Btbd11 does not outcompete TARP γ2 for Psd-95 binding slots and promotes protein complexes via phase separation

We show that increasing concentration of Btbd11 relative to Psd-95 and TARPγ2-CT does not prevent binding of TARPγ2 to Psd-95. By contrast, it promotes the formation of large phase separated condensates containing all three proteins. As both Btbd11 and TARPγ2 preferentially bind to Psd-95 PDZ2, though TARPγ2 also has high affinity for both PDZ 1 and ^3 30^, we originally hypothesized that Btbd11 might act as a competitor to TARPγ2 in a SynGAP-like manner. In ENs, SynGAP has been shown to phase separate with glutamatergic PSD proteins ^17^ and negatively regulates TARP binding to Psd-95 in an activity-dependent manner, occupying PDZ ‘slots’ and preventing TARPs from binding until CamKII-dependent phosphorylation of SynGAP leads to its dispersion from Psd-95 ^37,38^. However, high levels of Btbd11 do not diminish TARP binding to Psd-95, suggesting the proteins either have the capacity to bind at different PDZ domains, or multimerization of Psd-95 enables a complex to form even with both proteins binding to PDZ2 domains. This is in line with the structure and scaffolding nature of Psd-95; the three isolated PDZ domains coordinate multiple interactors, which promotes the ability to coordinate dynamic response to stimuli at the synapse ^51^. Taken together, the prediction that Btbd11 is likely a strong driver of phase separation, the presence of hallmark phase separation properties including FRAP ^52^ and puncta that coalesce over time, and that a higher concentration of Btbd11 can lead to increased size and inclusion of glutamatergic PSD molecules in condensates, suggest that Btbd11 contributes to IN-PSD organization and maintenance through by assembling dynamic formations of these critical proteins.

It is worth noting that the increased condensate size resulting from Btbd11 inclusion may be due to dimerization or multimerization of the proteins involved; Psd-95 is known to dimerize through its N-terminus ^28,53^ and Btbd11 is capable of multimerization (most clearly demonstrated by fibril formation when overexpressed^32^). Though the extent to which this is occurring at the synapse cannot be extrapolated from our data in HEK cells, our data suggest the possibility that Btbd11 supports Psd-95 multimerization via LLPS, supporting diverse protein-protein interactions within synapses.

### Synaptic localization and nanoscale architecture of Btbd11

Using a combination of confocal and STED microscopy, we examined the spatial localization of Btbd11 relative to Psd-95 at the synapse, finding a positive correlation between Psd-95 and Btbd11 synaptic intensity and NC number, but that Btbd11 and PSD-95 NCs were differentially distributed within the synapse and only colocalized incompletely within the synapse. The scaffolding complex of Psd-95 extends in a vertical, striated manner through the PSD that has been well documented; palmitoylated Psd-95 extends perpendicular to the membrane, where it binds to GKAP, Shank, and Homer to form a stable yet tunable platform via interactions with the actin cytoskeleton ^54^. We found that Btbd11 localizes beneath Psd-95 relative to the membrane, providing insight into how Btbd11 fits into the stratified PSD organization both overall and in relation to the important scaffolding of Psd-95. Given that the two proteins interact both through direct binding and phase condensation, it was particularly curious that the colocalization of the two was not more uniform. This is likely due to the fact that the antibody we used to visualize Btbd11 binds to an epitope within the N-terminus, and Btbd11 is a large protein (∼120kD). While Btbd11’s C-terminus can directly bind Psd-95, predictive modeling suggests the N-terminal portion is conformationally opposite and extends outwards, which is in line with our STED imaging (and may explain the separation of Btbd11 and Psd-95 signal, both at *en face* and on side synapses). The distribution of the disordered N-terminal tail deeper in the PSD may then support Btbd11’s role in promoting phase separation through multivalent protein-protein interactions.

### Btbd11 impacts protein stability in an expression- and phase separation-dependent manner

Using FRAP, we demonstrate that Btbd11 itself undergoes exchange at the synapse at a timescale similar to Psd-95, TARP, and the AMPAR subunit GluA1, a property that is absent in N-terminal Btbd11 mutants. This clearly demonstrates how the propensity of a synaptic protein to undergo LLPS can regulate its mobility, as demonstrated previously with SynGAP^55^. It is noteworthy that loss of the N-terminal region of Btbd11 does not impact synapse localization—just the mobility of Btbd11 at the synapse. Further, we show that Btbd11 overexpression impacts TARPγ2 and GluA1 stability at putative synapses, though strikingly, in opposite directions. Stabilization of TARPγ2 was expected following the findings that Btbd11 overexpression stabilizes Psd-95^32^ and Sap-102 (Fig 4), so the finding that Btbd11 overexpression led to an increase in the GluA1 mobile fraction was unexpected. This could be due to experimental differences; the use of SEP-GluA1 provided a readout of solely surface receptors while EGFP-TARPγ2 reports both surface and internalized protein. Differences in SEP and EGFP-tagged AMPAR recovery have been previously noted ^56^, in which the mobile surface fraction was greater when measured with SEP than EGFP (which was estimated at ∼20% internal receptors), though these experiments examined GluA2 in ENs. Further, though TARPγ2 is enriched in INs, AMPARs also interact with other TARP isoforms that influence surface and synapse targeting which could also contribute to this apparent difference ^31^. Reassuringly, the mobility of SEP-GluA1 in Btbd11 KO neurons showed a strong opposite effect to the overexpression data, highlighting a bidirectional effect. Crucially, overexpression of Btbd11 showed a significant difference compared to Btbd11ΔN, which itself was indistinguishable from control, suggesting the impact of Btbd11 on PSD protein stability is indeed dependent on its ability to undergo and promote phase separation. This is also underscored by the fact that Btbd11ΔN—which itself is very immobile at synapses—did not have any effect on the dynamics of synaptic proteins, which would have been expected if the stabilization effect was simply a result of protein-protein interactions (in the absence of LLPS).

AMPARs are highly dynamic and diffuse laterally following membrane insertion until capture at the synapse by scaffolding proteins, where they can then be immobilized in the PSD, unbind and diffuse away, or are internalized through endocytosis. One explanation for an increase in the GluA1 mobile fraction in Btbd11 overexpression is that Btbd11 may play a role in recruiting AMPARs to the phase separated PSD from the extra synaptic AMPAR pool. In an overexpression Btbd11 model, without additional MAGUK slots or activity-dependent changes occurring, Btbd11-recruited excess AMPARs would be unable to bind to scaffolding in the PSD and rapidly diffuse back out, leading to the observed enhancement of GluA1 mobility. Thus, it follows that a phase separation null-variant would have no impact on GluA1 mobility, as the loss of phase separation causes Btbd11 itself to remain in a more static state. A limitation of using FRAP is a lack of discrete data discriminating between different receptor pools; we cannot distinguish lateral diffusion of surface receptors freely diffusing in the membrane from recycling receptor pools delivered to the membrane through exocytosis. The surprising mismatch in measured TARPγ2 and GluA1 mobility following Btbd11 overexpression, given that nearly all AMPARs contain TARPγ2 ^31^, and vice versa, may be thus explained in the future by measuring different pools of surface versus internalized receptors or utilizing MINFLUX imaging for ultra-precise single-molecule localization tracking^57^.

### Synaptic PSD protein expression is reduced in Btbd11 KO cultured neurons

Following our functional FRAP findings that Btbd11 impacts PSD protein mobility, we next investigated how Btbd11 expression impacts PSD protein expression using immunofluorescence, finding a significant decrease in synaptic density and area or intensity in Btbd11 KO neurons compared to control. These results complement previous findings that Btbd11 KO mice displayed similarly reduced Psd-95 puncta area in parvalbumin positive INs in the visual cortex ^32^. Beyond Psd-95, we show a decrease in both TARPγ2 and GluA1 density, as measured by the number of puncta colocalized with presynaptic VGlut1, and decreased synaptic puncta intensity. These measures are consistent with the idea that loss of Btbd11 disrupts Psd-95 organization and abundance, which sequentially leads to a loss of GluA1 clustering via fewer available docking sites for TARPγ2 binding.

### Btbd11 KO impairs spontaneous calcium transients in neuronal cultures

The observed decrease in Psd-95, TARPγ2 and GluA1 following Btbd11 KO is consistent with impaired glutamatergic recruitment of INs previously reported ^32^. To confirm whether this was the case, we investigated synapse function in INs in control and Btbd11 KO neurons using calcium imaging. We saw a significant decrease in calcium transients in Btbd11 KO neurons compared to control, providing strong evidence that the postsynaptic disruptions we have observed manifest in quantifiable functional deficits in glutamatergic transmission. To focus primarily on spontaneous calcium transients originating from AMPARs, we pharmacologically blocked NMDARs. AMPARs in interneurons are typically characterized by an absence of the calcium-impermeable GluA2 subunit ^7^, and as such are highly permeable to calcium influx ^1,2^. Although we saw a decrease in AMPAR expression in KO neurons, the vast decrease in frequency and amplitude of calcium transients in the KO condition were even more dramatic than expected: multiple KO cells did not experience a single event. This disproportionate functional loss could reflect disrupted Psd-95 engagement with neuroligins and their presynaptic neurexin binding partners^58^, perturbing the retrograde trans-synaptic signaling that coordinates release with postsynaptic organization^59^. Though no difference in presynaptic release probability was previously noted in Btbd11 KO mice cortex^32^, GluSnFr imaging could provide insight into any alterations to presynaptic glutamate *in vitro*. Collectively, these data demonstrate that Btbd11 is not only a critical determinant of glutamatergic synapse organization in INs, but is required for glutamatergic transmission at these synapses.

### Btbd11 KO disrupts Psd-95 nanoscale organization

We used super resolution imaging to quantify changes in Psd-95 NC organization and observed decreased NC number and size in Btbd11 KO neurons compared to control. These data provide a key insight into how Btbd11 supports glutamatergic transmission in INs; smaller and fewer NCs impact the organization and efficiency of receptor and signaling complexes^18,60,43^. This is the first demonstration of a molecular mechanism that regulates Psd-95 NCs in INs. Notably, while the reduction in NC number likely reflects the smaller synapse area, with which NC number positively correlates in both genotypes, the reduction in NC size cannot be attributed to synapse scaling alone and instead points to a specific disruption of nanoscale PSD organization. Consistent with this, KO neurons show a significantly reduced PSD width and area, with no change in Psd-95 membrane distribution. Thus, even though AMPARs and Psd-95 remain at the PSD in the absence of Btbd11, receptors may be insufficiently concentrated within NCs and misaligned from their presynaptic release sites, a disorganization that could manifest as dramatically reduced glutamatergic signaling and warrants direct investigation in future studies.

Though general qualities are similar between EN and IN nanostructure, notable differences have been documented ^6^. Previous work has shown that IN-PSDs are relatively larger and contain the same overall number of NCs compared to EN-PSDs, while IN-PSD NCs tend to be larger and denser. These data suggest IN-PSDs contain more abundant space between Psd-95 NCs in which proteins such as AMPARs must diffuse between before capture and stabilization^6^. Further, IN-PSDs display a stronger trans-synaptic alignment between Psd-95 and presynaptic protein Munc13-1, which regulates docking and priming of synaptic vesicles at the active zone and whose NCs are thought to mark release sites^61–63^, than EN-PSDs. This enhanced alignment of presynaptic release and postsynaptic receptor sites in IN-PSDs suggests a stronger excitatory drive, possibly requiring specialized cell-type specific machinery such as Btbd11. Taken together with our current findings that Btbd11 expression regulates both MAGUK/TARPγ2/AMPAR mobility and Psd-95 NC organization, Btbd11 may be uniquely positioned to support glutamatergic transmission by promoting TARP/AMPAR recruitment and shuttling receptors through the phase-separated PSD to Psd-95 NCs.

## Materials and Methods

### Animal care

All animals were treated in accordance with the Tufts University Animal Care and Use Committee guidelines. Transgenic mice and timed pregnant Sprague-Dawley rats were used in this study to generate primary neuronal cultures. All animals were housed in a temperature- and humidity-controlled vivarium with unlimited access to drinking water and food.

### Cell Culture

#### HEK cells

HEK cells were grown on 6-well plates (for biochemistry) or on collagen (Advanced Biomatrix) coated glass coverslips in 12-well plates (for immunofluorescence and live-cell imaging) in DMEM (Gibco) supplemented with 10% Fetal Bovine Serum (ThermoFisher) and Penicillin-Streptomycin antibiotics (100 U/ml, Thermo Fisher Scientific). Cells were maintained at 37C with 5% CO_2_ and used up to Passage 20 (P20). HEK cells were transfected via Calcium Phosphate precipitation for between 4-5 hours followed by 100% media change the day prior to collection (for biochemistry) or fixation (immunofluorescence). For live-imaging experiments, cells were transfected with Lipofectamine 2000 (Invitrogen) for one hour followed by 100% media change and imaged 4-6 hours later.

#### Primary cultured rat neurons

Timed pregnant Sprague-Dawley rats were purchased (Charles River) and dissected at embryonic day 18. Hippocampi of 4-5 embryos were dissected into ice-cold dissection media (10x HBSS (Gibco), Penicillin-Streptomycin (Gibco), sodium pyruvate (Gibco), 10mM HEPES (Gibco), 30mM Glucose (Sigma) and Milli-Q water) prior to chemical dissociation via papain (Worthington) and subsequent trituration via P1000 pipette. Cells were plated at 100,000/well onto glass coverslips coated with 1mg/mL Poly-L-Lysine (Sigma) of a 12-well plate (for immunofluorescence and GCaMP live imaging) and 350,000 per glass-bottom 35mm dish (for live-imaging) coated with Poly-L-Lysine. Neurons were grown in Neurobasal media (Gibco) supplemented with Penicillin-Streptomycin (Gibco), 2mM Glutamax (Gibco), 5% horse serum (Gibco), and 2% B27 (Gibco). A full media change was performed two hours after plating cells. One day after plating cells (DIV1), 100% of media in 35mm dishes replaced with clear serum-free Neurobasal, while 70% of media in 12-well plates was replaced with serum-free Neurobasal media. Cells were fed once a week and maintained at 37C with 5% CO2. For Btbd11/ΔN overexpression FRAP, neurons were transfected at DIV12 with 9uL Lipofectamine 2000 per dish mixed with plasmid DNA (3µg mDlx-azurite, 0.5µg CAG-mCherry-Btbd11 or CAG-mCherry-Btbd11ΔN, and either 0.5µg CAG-GFP-TARPγ2 or 1µg mDlx-Sep-GluA1) for 45 minutes followed by a full media change containing 50% media retained prior to transfection and 50% freshly prepared Neurobasal. Neurons for Btbd11 KO FRAP were generated by transfecting cells at div4 under the same conditions as overexpression other than reducing plasmid concentrations (0.5µg mDlx-azurite, 0.25µg mDlx-SEP-GluA1, including 1µg CRISPR guide in KO condition only). For immunofluorescence, neurons were fixed at DIV14 or 15 with 4% PFA (Electron Microscopy Science) for 15 minutes or, if staining for Btbd11, 100% methanol for 20 minutes at -20C.

#### Primary cultured mouse neurons

P0 Btbd11^Fl/Fl^ pups were dissected and plated as described above for rat neurons. Hippocampal neurons were plated onto Poly-L-Lysine coated (1 mg/ml) glass coverslips at 150,000/well for 12-well plates, followed by a 100% media change after 2 hours. 70% of media was replaced with serum-free media one day after plating cells, then fed once per week. Cells were maintained at 37C with 5% CO2. To generate control and KO cells, 2 µl of AAVs were added across 6 wells of a 12-well plate at div1. pAAV.CMV.HI.eGFP-Cre.WPRE.SV40, serotype AAV9, titer 1×10¹³ vg/mL (a gift from James M. Wilson, Addgene plasmid #105545) and AAV.CMV.PI.EGFP.WPRE.bGH, serotype AAV9; 1×10¹³ vg/mL (a gift from James M. Wilson, Addgene plasmid #105530) were purchased from Addgene. For immunofluorescence, neurons were fixed at div14 or 15 with 4% PFA (Electron Microscopy Science) for 15 minutes. For GCaMP imaging, control and KO neurons were generated via pAAV-Ef1a-mCherry, serotype AAV8, titer 1×10¹³ vg/mL (a gift from Karl Deisseroth, Addgene plasmid #114470) or pAAV-Ef1a-mCherry-IRES-Cre, serotype AAV8, titer 1×10¹³ vg/mL (a gift from Karl Deisseroth, Addgene plasmid #55632) delivery at div1. AAV-mDlx-jGCaMP8s-WPRE, serotype AAV9, titer 1×10¹³ vg/mL (a gift from Loren Looger, Addgene plasmid #176755) was delivered to both control and KO cells at div7.

### Cloning and molecular biology

All constructs were generated using HiFi assembly (New England Biolabs). Psd-95-mCherry PDZ1 and Psd-95-mCherry PDZ2 mutants were generated by cutting and cloning mutated and wildtype PDZ1 and PDZ2 into a pCAG-Psd-95 backbone, taken from existing Psd-95-mCherry PDZ1,2 mutants^32^. Psd-95-azurite was generated by replacing mCherry in the aforementioned plasmid. The sequence of all constructs was confirmed with DNA sequencing prior to use. GFP-TARPγ2-CT was a gift from Richard Huganir. The Btbd11 CRISPR knockdown plasmid pU6-(BbsI) CBh-Cas9-T2A-mCherry was a gift from Ralf Kuehn (Addgene plasmid #64324). The guide sequence target was 5-CGAAGCGCCCAAATTCACCG-3’.

### GST-Pulldowns

For all GST pulldowns, HEK cells were grown in 6-well plates to ∼70% confluence and transfected with DNA plasmids using calcium phosphate. 1µg of each DNA plasmid was transfected per well, except for competition experiments in Figure 2C, in which the molar equivalent to 1µg of PSD-95 of Btbd11 and TARP were transfected. The following day, cells were rinsed once with 1X PBS prior to lysis with ice-cold lysis buffer (1X Tris-Buffered Saline, 0.5% NP-40, 10mM NaPPi, 10mM NaF, 1mM Na_3_VO_4_, and Roche protease/phosphatase inhibitor.) Cells were solubilized by rotation for 20 min at 4C, then centrifuged at 17,000g for 15 min to remove cellular and nuclear debris. A small percentage of supernatant was retained to run as input, while the rest was incubated for two hours at 4C with 30mL of pre-washed Glutathione Sepharose 4B bead slurry (GE Healthcare). Beads were then washed in spin columns (Corning) 5 times with 500uL lysis buffer per wash, then eluted with 30uL of 2X Laemmli Buffer (Sigma) at 70C for 3-5 minutes. Samples were either stored in -20 or run the same day on western blot.

### Western Blotting

Samples were heated at 70C for 15 minutes prior to running blots. Precast BIS-TRIS 4-12% or 8% gels (ThermoFisher) were used, depending on target protein molecular weight, with NuPage MOPS running buffer (ThermoFisher). Gels were run for 45 minutes at 200V before transfer to nitrocellulose membrane (ThermoFisher) for 1 hour at 100V. Membranes were blocked in 3% BSA (Sigma) in 0.1% Tween-20 (Sigma Aldritch) in Tris-Buffered Saline (Thermo Fisher Scientific) (TBST) for 1 hour before incubation with primary antibodies for 1-2 hours at RT or overnight at 4C. Primary antibodies (listed below) were made up in 0.1% TBST with 3% BSA. Blots were washed 5-7 times with 0.1% TBST prior to secondary incubation, also in 0.1% TBST for 1-2 hours at RT, and all secondaries (Li-Cor) were fluorescently conjugated with either 680 or 800 dyes and used at 1:10000 concentration. The membrane was washed 5-7 times with 0.1% TBST prior to imaging with Li-Cor Odyssey DLx near-infrared scanner. Bands were quantified in EmpiriaStudio (Li-Cor). Primary antibodies used: homemade anti-Btbd11 (rabbit polyclonal, 1:1000) ^32^ anti-mCherry (abcam AB125096 IgG2_A_) was used at 1:5000, mouse anti-GFP (Santa Cruz SC-9996 IgG2_A_,) was used at 1:2000, mouse anti-GluA1 (4.9D IgG2_A_; a gift from Richard Huganir) was used at 1:5000.

### FRAP

All FRAP experiments were performed using an oil-immersive 63x objective on a Zeiss confocal microscope (LSM980). HEK cells were transfected using Lipofectamine 4-6 hours prior to imaging, and coverslips were moved from DMEM to a custom 3D-printed imaging chamber containing aCSF (120mM NaCl, 5 mM KCl, 10mM HEPES, 10mM glucose, 2mM CaCl_2_ and 1mM MgCl_2_). Cells were live-imaged in a humidity-controlled chamber maintained at 37C, with the chamber containing 5% CO2 for imaging of neurons in media or without CO2 for imaging of HEK cells or GCamp neurons in aCSF, which did not require CO2 due to the presence of HEPES. Interneurons were identified via mDlx-azurite signal, and a high-resolution z-stack was taken of each cell prior to bleaching. A three-slice Z-stack at 0.5-micron steps were taken at each timepoint, and two timepoints were acquired at baseline prior to bleaching. Bleaching was performed by laser at 80-100% intensity for 15 iterations. For TARPγ2 and GluA1 FRAP, each timepoint was imaged 60 seconds apart for 30 minutes. SAP102 FRAP timepoint was every 30 seconds for 5 minutes. For each experiment, plasmid concentrations, laser power, and number of bleaching repetitions was optimized, then kept consistent. For HEK cell FRAP, 0.5 µg azurite-Psd-95 was transfected with the molar equivalent of mCherry-Btbd11 and GFP-TARPγ2-CT (0.6477 and 0.4311 µg, respectively). For Btbd11 and Btbd11ΔN FRAP in neurons, 0.4 µg of either Btbd11 plasmid was transfected with 1 µg GFP-FingR-PSD-95 and 3 µg mDlx-azurite. For GFP-TARPγ2 FRAP in neurons, 0.5 µg of GFP-TARPγ2 was transfected with 3 µg of mDlx-azurite, alone or with 0.5 µg of either Btbd11 plasmid. For GluA1 FRAP with Btbd11 overexpression, 1 µg of mDlx-SEP-GluA1 was transfected with 3 µg of mDlx-azurite, alone or with 0.5 µg of either Btbd11 plasmid. For GluA1 FRAP with Btbd11 KO, 0.25 µg of GFP-TARPγ2 was transfected with 1 µg of mDlx-azurite, alone or with 1 µg of Btbd11 KO CRISPR guide.

### Immunocytochemistry

Prior to fixation, cells were washed once with sterile room temperature phosphate-buffered saline (PBS). For most experiments, cells were then fixed with 4% paraformaldehyde (Electron Microscopy Science) and 4% sucrose (Sigma) in PBS for 15 minutes at room temperature. For experiments staining for endogenous Btbd11, cells were instead subject to a 100% ice-cold methanol fixation at - 20C for 20 minutes. Following fixation, cells were washed three times with PBS to remove any trace of fixative.

For Psd-95 staining, all steps were performed on the benchtop at room temperature. Neurons were first washed 2-3 three times with PBS prior to permeabilization with 0.1% Triton-X and 10% Normal Goat Serum (VectorLabs) in PBS for 15 minutes. Neurons were briefly washed once with PBS before primaries were added in 0.1% Triton-X and 3% Normal Goat Serum (VectorLabs) in PBS for 1.5 hours. FluoTag-X2 anti-PSD-95 (NanoTag Biotechnology #N3702) was pre-conjugated to abberior STAR 635P and used at 1:500 for confocal imaging as in Figure 5A, while 1:250 concentration was used for STED in Figure 3. Mouse IgG2A GAD67 (Millipore Sigma #MAB5406) was used at 1:500, and either guinea pig vGlut1 (Synaptic Systems #135304) was used at 1:1000 (for confocal imaging) or guinea pig Bassoon (Synaptic Systems #141 318) were used at 1:5000 (for STED imaging). For endogenous Btbd11 staining (Figure 3A), rabbit Btbd11 (abclonal # A20192) was used at 1:1000. Cells were then washed three times with PBS before secondaries were added in 0.1% Triton-X and 3% Normal Goat Serum (VectorLabs) in PBS for one hour. Secondaries were all used at a 1:1000 concentration. Goat anti-mouse IgG2A DyLight 405-conjµgated secondary (Jackson Immuno Research #115-477-186) was used against GAD67, and goat anti-guinea pig Alexa Fluor 488-conjµgated secondary (ThermoFisher #A11073) was used against VGlut1 or Bassoon. Following secondary incubation, cells were washed three times in PBS followed by a single rinse of the coverslip in distilled water prior to mounting on a SuperFrost Plus microscope slide (Fisher Scientific #12-550-15) with ProLong Glass Antifade mounting media (ThermoFisher #P36984) or abberior Solid Antifade media for STED samples (abberior #DAD1494905ABB). Slides were maintained in the dark at 4C.

For TARPγ2 and GluA1 staining in Figure 5B-C, after fixation neurons were first washed 2-3 three times with PBS before a 20-minute permeabilization with 0.2% Triton-X in PBS. Neurons were then washed another 3 times with PBS before blocking in 10% Normal Goat Serum (VectorLabs) and 5% Bovine Serum Albumin (Sigma-Aldritch) in PBS for an hour. Primary antibodies were added in the same blocking buffer and left overnight at 4C. Rabbit TARPγ2 (Millipore Sigma #AB9876) or mouse IgG2_B_ GluA1 (clone 11b8; a gift from Eric Gouaux) was used at 1:500 and 1:1000 respectively, mouse IgG2A GAD67 (Millipore Sigma #MAB5406) was used at 1:500, and guinea pig VGlut1 (Synaptic Systems #135304) was used at 1:1000. The next day, cells were washed 3 times with PBS before adding secondaries for 1.5 hours at room temperature, in the same buffer as primary and blocking. Secondaries were all used at a 1:1000 concentration. Goat anti-rabbit Alexa Fluor 647-conjµgated secondary (ThermoFisher #A21244) was used against TARPγ2, goat anti-IgG2B Alexa Fluor 488-conjugated secondary (#A21131) was used against GluA1, goat anti-mouse IgG2A Alexa Fluor 594- or 488-conjµgated secondary (ThermoFisher #A21134 and #A21131, respectively) was used against GAD67, and goat anti-guinea pig DyLight 405-conjµgated secondary (Jackson Immuno Research #106-475-003) was used against VGlut1. Following secondary incubation, cells were washed three times in PBS followed by a single rinse of the coverslip in distilled water prior to mounting on a SuperFrost Plus microscope slide (Fisher Scientific #12-550-15) with ProLong Glass Antifade mounting media (ThermoFisher #P36984). Slides were maintained in the dark at 4C.

Image acquisition was carried out on an oil-immersive 63x objective on a Zeiss confocal microscope (LSM980). Z-stacks of approximately 5-6µm at 0.5µm steps were acquired, and maximum projections of stacks were used for all analysis.

### STED

STED images were acquired on an Abberior Facility Line STED microscope equipped with an Olympus IX83 base and UPlanXApo 60x/1.42 NA oil immersion objective, with pulsed 405, 488, 561, and 640 nm excitation lasers and a pulsed 775 nm STED depletion laser. All imaging was performed at least 1 hour after starting up the microscope and lasers to stabilize the system, and the microscope was calibrated approximately every 4-6 hours. We identified GAD67+ interneurons using the 405 channel by eyepiece, then acquired 1- or 2-color STED images with confocal presynaptic bassoon for KO and endogenous Btbd11, respectively. Three 15-20µm sections of dendrite per neuron were imaged, selected from six neurons across two replicate coverslips per batch, from three independent batches of neurons per condition.

For endogenous Btbd11 and Psd-95 imaging, Abberior STAR635p dye (Psd-95) was excited by the 640 nm laser at 25% power and depleted by the STED laser at 25% power, with accumulation of 7 lines. Alexa Fluor 594 (Btbd11) was excited by the 561 nm laser at 60% power and depleted by the STED laser at 30% power, with accumulation of 7 lines. For Btbd11 KO, Abberior STAR635p dye (Psd-95) was excited by the 640 nm laser at 30% power and depleted by the STED laser at 25% power, with accumulation of 7 lines. Corresponding confocal-resolution images were acquired alongside each STED image.

### GCaMP imaging

All GCaMP imaging was carried out using an oil-immersive 63x objective on a Zeiss confocal microscope (LSM980) at 2X zoom. div14-17 neurons were live-imaged in a humidity-controlled chamber maintained at 37C in aCSF (120mM NaCl, 5 mM KCl, 10mM HEPES, 10mM glucose, 2mM CaCl_2_ and 1mM MgCl_2,_ pH 7.30). aCSF was further supplemented with 1µM TTX (Tocris) and 1M Mg2+ (Invitrogen, #AM9530G). Following baseline imaging, 50 mM AP5 was applied to block NMDAR and preferentially capture spontaneous calcium transients originating from AMPARs. Viral mCherry expression was visually confirmed in each neuron prior to imaging. Neurons were imaged in 30-second intervals at 200 frames per second using camera streaming. Finally, 20µM DNQX was applied to block AMPARs, confirming calcium signals originated from AMPARs. GCaMP interneuron expression specificity under the mDlx promoter was validated using GAD67 staining (20 out of 20 mDlx cells were GAD67-positive, data not shown).

### Analysis

#### FRAP

For HEK cell FRAP experiments, 1-2 condensates or aggregates were bleached per cell, with another ROI drawn around unbleached protein and a background ROI. ROI intensities were measured at each timepoint. Background signal was subtracted from the puncta signal, which was then normalized to the average intensity of the unbleached puncta at each time point (to account for the low levels of acquisition bleach over time). The signal in the bleached ROIs was then normalized such that the average baseline was centered on 1 and the post-bleach time point was 0. For each condensate or aggregate, the estimated plateau (maximum recovery) was estimated with a one phase exponential fit (in GraphPad Prism 9).

For all neuronal FRAP experiments, 7 putative synaptic puncta of the protein of interest were bleached, 3 were selected as representative unbleached, and 3 ROIs of similar size were also selected in the background. Files were run through a custom Python script to split channels and create a new file with only the channel of the protein of interest and a generic name to blind the person analyzing each file to the condition. In Fiji, maximum image projections were generated for the timeseries. A 1-pixel median filter and a rigid-body transformation was applied to correct for drift (MultiStackReg plugin). Puncta were analyzed only if bleached to below 30% of the average baseline intensity measurements, and unbleached puncta were analyzed if the timepoint immediately following the bleach event did not deplete below 80% of the average baseline intensity. Background signal was subtracted from the puncta signal, which was then normalized to the average intensity of the unbleached puncta at each time point (to account for the low levels of acquisition bleach over time). The signal in the bleached ROIs was then normalized such that the average baseline was centered on 1 and the post-bleach time point was 0. Puncta were averaged per neuron and only used if 3 or more puncta fit the bleaching criteria. For each cell, the estimated plateau (maximum recovery) was estimated with a one phase exponential fit (in GraphPad Prism 9).

#### STED

Btbd11 and Psd-95 (Alexa-594 and STAR635P channels, respectively) were deconvolved from raw STED images in Abberior LightBox. All data were analyzed using custom FIJI macros and Python. Dendrite ROIs were manually drawn in each image to segment IN dendrites prior to synaptic detection via unsharp masking. To remove background and identify synapses, deconvolved Psd-95 STED images filtered by gaussian blur with a radius of 30 pixels were subtracted from STED images filtered by gaussian blur with a radius of 3 pixels, as previously described ^42^. Detections were binarized via Otsu threshold and objects filtered with an area cutoff of 0.01µm^2^. ROIs were further filtered to only include PSDs that contained positive Bassoon signal. Btbd11 STED images were detected similarly, with additional filtering to only include Btbd11 detections within Psd-95 and Bassoon-positive ROIs.

Psd-95 NC ROIs were detected with a low (0) and high (3) gaussian blur with an area cutoff of 0.001µm^2^ and watershed segmentation and filtered to only include NCs within defined Bassoon-positive Psd-95 synapse ROIs. Btbd11 NC ROIs were detected with low (0) and high (4) gaussian blur, but otherwise identical to Psd-95 detection. The obtained Psd-95 and Btbd11 NC masks were used to measure correlation and colocalization between Psd-95 and Btbd11 NC number and size per synapse. Colocalization was defined as any intersection of union (IoU) value > 0 for NC shapes of each protein, generated via unsharp thresholding.

On side synapses were manually selected and exported as individual synapses. Psd-95 and Bassoon ROIs were generated again using the unsharp detection method on individual channels with a gaussian blur with a radius of 30 pixels subtracted from images filtered by gaussian blur with a radius of 3 pixels and Otsu thresholding. To generate line scans, the Feret diameter was produced for Psd-95 channel. A line of the same width as the Feret line was created perpendicular and centered to run through the centroid of the combined Psd-95-Bassoon shape, with an excess of 5 pixels in each direction. Distance and intensity for Psd-95, Btbd11, and Bassoon were calculated at each point along the line. The maximum intensity for each channel was normalized between 1 and 0. Each line scan was rescaled, so the Psd-95 maximum peak was set to distance 0 to generate relative distance of each channel to Psd-95. Each Psd-95 peak relative to Bassoon was scaled in the same direction so Psd-95 was always at the beginning of the line scan, with Btbd11 relative to Psd-95. The resulting maximum peak of each channel per synapse was then used to generate an average line scan plot and measure Btbd11 peak offset relative to Psd-95.

Bassoon-positive Psd-95 synapses in Btbd11 KO neurons were analyzed in an identical manner as above. Resulting Psd-95 NC number and size was compared between Control and KO for en face synapses, while the difference in average Feret diameter was measured between Control and KO for on side synapses. Psd-95 thickness, or depth into the PSD away from the membrane, was calculated using the distance between the outermost line scan points where the signal reached ≥10% of its peak.

### GCaMP

To quantify dendritic calcium activity, 3-5 sections of individual dendrites per neuron were first manually segmented from time-series imaging data and isolated using binary masks. For each dendrite, the two-dimensional fluorescence signal was reduced to a one-dimensional representation by averaging pixel intensities along the transverse axis (width) of the dendrite, yielding a position-by-time matrix of fluorescence values along the dendrite. Changes in fluorescence were expressed as ΔF/F₀, where F₀ was defined using a moving percentile baseline to account for slow baseline drift and photobleaching over time. Specifically, for each dendritic position, F₀ was computed as the lower 20th percentile within a sliding temporal window of 250 frames. Calcium events were identified by visualizing the resulting ΔF/F₀ position over time heatmaps and manually annotating ROIs spanning both spatial and temporal dimensions. Within each ROI, the position of the maximal ΔF/F₀ value was defined as the site of peak activity and taken as a proxy for the synaptic locus. The peak ΔF/F₀ at this position was used as a measure of event amplitude.

### Quantification and Statistical Analysis

GraphPad Prism 9 was employed for all statistical analysis and most graphs. Figures were plotted in GraphPad Prism 9 or using Python and Jupyter Notebooks. All FRAP, STED, immunofluorescence, and GCaMP analyses were performed on blinded files to prevent bias. All data were tested for normality prior to statistical testing, and data with two comparison groups that passed normality tests were further tested for statistical differences with two tailed unpaired t-test if variance was equal, or Welch’s t-test for data with unequal variance. Data with more than two comparison groups that passed normality tests were tested for statistical differences using one-way ANOVA and post hoc Dunnett’s multiple comparison tests. Data that did not pass normality tests were subject to comparison with Mann-Whitney tests, Kruskal-Wallis test for multiple unmatched comparison groups, or Friedman test for matched comparisons. All statistical tests shown in Supplementary Table 1.

## Acknowledgments

We would like to thank members of the Bygrave and Blanpied laboratories for helpful discussions, and Dr. Shilpa Dilip Kumar for assistance on STED imaging. This work was supported by F31MH140553 (M.B.B.), R21MH141559 (A.D.L. and T.A.B), R37MH008046 (T.A.B.), and R00MH124920 (A.M.B). STED images were acquired in the University of Maryland School of Medicine Center for Innovative Biomedical Resources Core Confocal Facility, with support from the University of Maryland Medicine Institute for Neuroscience Discovery (UM-MIND).

## References

1. Geiger, J. R. P. et al. Relative abundance of subunit mRNAs determines gating and Ca2+ permeability of AMPA receptors in principal neurons and interneurons in rat CNS. Neuron 15, 193–204 (1995).

2. Geiger, J. R. P., Lübke, J., Roth, A., Frotscher, M. & Jonas, P. Submillisecond AMPA Receptor-Mediated Signaling at a Principal Neuron–Interneuron Synapse. Neuron 18, 1009–1023 (1997).

3. NyÍri, G., Stephenson, F. A., Freund, T. F. & Somogyi, P. Large variability in synaptic n-methyl-d-aspartate receptor density on interneurons and a comparison with pyramidal-cell spines in the rat hippocampus. Neuroscience 119, 347–363 (2003).

4. Kawaguchi, Y., Karube, F. & Kubota, Y. Dendritic Branch Typing and Spine Expression Patterns in Cortical Nonpyramidal Cells. Cerebral Cortex 16, 696–711 (2006).

5. Chang, M. C. et al. Narp regulates homeostatic scaling of excitatory synapses on parvalbumin-expressing interneurons. Nat Neurosci 13, 1090–1097 (2010).

6. Dharmasri, P. A., Levy, A. D. & Blanpied, T. A. Differential nanoscale organization of excitatory synapses onto excitatory vs. inhibitory neurons. Proc. Natl. Acad. Sci. U.S.A. 121, e2315379121 (2024).

7. Hong, I. et al. Calcium-permeable AMPA receptors govern PV neuron feature selectivity. Nature 635, 398–405 (2024).

8. Hu, H., Gan, J. & Jonas, P. Fast-spiking, parvalbumin ^+^ GABAergic interneurons: From cellular design to microcircuit function. Science 345, 1255263 (2014).

9. Chung, D. W., Fish, K. N. & Lewis, D. A. Pathological Basis for Deficient Excitatory Drive to Cortical Parvalbumin Interneurons in Schizophrenia. AJP 173, 1131–1139 (2016).

10. Lewis, D. A., Curley, A. A., Glausier, J. R. & Volk, D. W. Cortical parvalbumin interneurons and cognitive dysfunction in schizophrenia. Trends in Neurosciences 35, 57–67 (2012).

11. Nahar, L., Delacroix, B. M. & Nam, H. W. The Role of Parvalbumin Interneurons in Neurotransmitter Balance and Neurological Disease. Front. Psychiatry 12, 679960 (2021).

12. Mukherjee, A., Carvalho, F., Eliez, S. & Caroni, P. Long-Lasting Rescue of Network and Cognitive Dysfunction in a Genetic Schizophrenia Model. Cell 178, 1387–1402.e14 (2019).

13. Hosokawa, T. & Liu, P.-W. Regulation of the Stability and Localization of Post-synaptic Membrane Proteins by Liquid-Liquid Phase Separation. Front. Physiol. 12, 795757 (2021).

14. Zeng, M. et al. Phase Transition in Postsynaptic Densities Underlies Formation of Synaptic Complexes and Synaptic Plasticity. Cell 166, 1163–1175.e12 (2016).

15. Chen, S. et al. Native postsynaptic density is a functional condensate formed via phase separation. Cell Reports 45, 116723 (2026).

16. Chen, X., Wu, X., Wu, H. & Zhang, M. Phase separation at the synapse. Nat Neurosci 23, 301–310 (2020).

17. Zeng, M., Bai, G. & Zhang, M. Anchoring high concentrations of SynGAP at postsynaptic densities via liquid-liquid phase separation. Small GTPases 1–9 (2017) doi:10.1080/21541248.2017.1320350.

18. Nair, D. et al. Super-Resolution Imaging Reveals That AMPA Receptors Inside Synapses Are Dynamically Organized in Nanodomains Regulated by PSD95. Journal of Neuroscience 33, 13204–13224 (2013).

19. Goncalves, J. et al. Nanoscale co-organization and coactivation of AMPAR, NMDAR, and mGluR at excitatory synapses. Proc. Natl. Acad. Sci. U.S.A. 117, 14503–14511 (2020).

20. Liu, P.-W., Hosokawa, T. & Hayashi, Y. Regulation of synaptic nanodomain by liquid–liquid phase separation: A novel mechanism of synaptic plasticity. Current Opinion in Neurobiology 69, 84–92 (2021).

21. Sun, S.-Y. et al. Correlative Assembly of Subsynaptic Nanoscale Organizations During Development. Front. Synaptic Neurosci. 14, 748184 (2022).

22. Schnell, E. et al. Direct interactions between PSD-95 and stargazin control synaptic AMPA receptor number. Proc. Natl. Acad. Sci. U.S.A. 99, 13902–13907 (2002).

23. Dakoji, S., Tomita, S., Karimzadegan, S., Nicoll, R. A. & Bredt, D. S. Interaction of transmembrane AMPA receptor regulatory proteins with multiple membrane associated guanylate kinases. Neuropharmacology 45, 849–856 (2003).

24. Ehrlich, I. & Malinow, R. Postsynaptic Density 95 controls AMPA Receptor Incorporation during Long-Term Potentiation and Experience-Driven Synaptic Plasticity. J. Neurosci. 24, 916–927 (2004).

25. Chen, X. et al. PSD-95 family MAGUKs are essential for anchoring AMPA and NMDA receptor complexes at the postsynaptic density. Proc. Natl. Acad. Sci. U.S.A. 112, (2015).

26. Chetkovich, D. M., Chen, L., Stocker, T. J., Nicoll, R. A. & Bredt, D. S. Phosphorylation of the Postsynaptic Density-95 (PSD-95)/Discs Large/Zona Occludens-1 Binding Site of Stargazin Regulates Binding to PSD-95 and Synaptic Targeting of AMPA Receptors. J. Neurosci. 22, 5791–5796 (2002).

27. Tomita, S., Fukata, M., Nicoll, R. A. & Bredt, D. S. Dynamic Interaction of Stargazin-like TARPs with Cycling AMPA Receptors at Synapses. Science 303, 1508–1511 (2004).

28. Xu, W. et al. Molecular Dissociation of the Role of PSD-95 in Regulating Synaptic Strength and LTD. Neuron 57, 248–262 (2008).

29. Opazo, P., Sainlos, M. & Choquet, D. Regulation of AMPA receptor surface diffusion by PSD-95 slots. Current Opinion in Neurobiology 22, 453–460 (2012).

30. Hafner, A.-S. et al. Lengthening of the Stargazin Cytoplasmic Tail Increases Synaptic Transmission by Promoting Interaction to Deeper Domains of PSD-95. Neuron 86, 475–489 (2015).

31. Yamasaki, M. et al. TARP γ-2 and γ-8 Differentially Control AMPAR Density Across Schaffer Collateral/Commissural Synapses in the Hippocampal CA1 Area. J. Neurosci. 36, 4296–4312 (2016).

32. Bygrave, A. M. et al. Btbd11 supports cell-type-specific synaptic function. Cell Reports 42, 112591 (2023).

33. Uversky, V. N. Protein intrinsic disorder-based liquid–liquid phase transitions in biological systems: Complex coacervates and membrane-less organelles. Advances in Colloid and Interface Science 239, 97–114 (2017).

34. Abramson, J. et al. Accurate structure prediction of biomolecular interactions with AlphaFold 3. Nature 630, 493–500 (2024).

35. Imamura, F., Maeda, S., Doi, T. & Fujiyoshi, Y. Ligand Binding of the Second PDZ Domain Regulates Clustering of PSD-95 with the Kv1.4 Potassium Channel. Journal of Biological Chemistry 277, 3640–3646 (2002).

36. Fernández, E. et al. Targeted tandem affinity purification of PSD-95 recovers core postsynaptic complexes and schizophrenia susceptibility proteins. Molecular Systems Biology 5, 269 (2009).

37. Araki, Y., Zeng, M., Zhang, M. & Huganir, R. L. Rapid Dispersion of SynGAP from Synaptic Spines Triggers AMPA Receptor Insertion and Spine Enlargement during LTP. Neuron 85, 173–189 (2015).

38. Araki, Y. et al. SynGAP regulates synaptic plasticity and cognition independently of its catalytic activity. Science 383, eadk1291 (2024).

39. Zeng, M. et al. Phase Separation-Mediated TARP/MAGUK Complex Condensation and AMPA Receptor Synaptic Transmission. Neuron 104, 529–543.e6 (2019).

40. Vendruscolo, M. & Fuxreiter, M. FuzDrop: sequence-based prediction of the propensity of proteins for liquid–liquid phase separation and aggregation. Nat Protoc 10.1038/s41596-025-01267-0 (2026) doi:10.1038/s41596-025-01267-0.

41. Anderson, M. C. et al. Trans-synaptic molecular context of NMDA receptor nanodomains. Nat Commun 16, 7460 (2025).

42. Sakamoto, H. et al. Synapse type-specific molecular nanoconfigurations of the presynaptic active zone in the hippocampus identified by systematic nanoscopy. Preprint at 10.1101/2022.03.11.483942 (2022).

43. Broadhead, M. J. et al. PSD95 nanoclusters are postsynaptic building blocks in hippocampus circuits. Sci Rep 6, 24626 (2016).

44. Dani, A., Huang, B., Bergan, J., Dulac, C. & Zhuang, X. Superresolution Imaging of Chemical Synapses in the Brain. Neuron 68, 843–856 (2010).

45. Akter, Y. et al. Combining nanobody labeling with STED microscopy reveals input-specific and layer-specific organization of neocortical synapses. PLoS Biol 23, e3002649 (2025).

46. Bensussen, S. et al. A Viral Toolbox of Genetically Encoded Fluorescent Synaptic Tags. iScience 23, 101330 (2020).

47. Zheng, C.-Y., Petralia, R. S., Wang, Y.-X., Kachar, B. & Wenthold, R. J. SAP102 Is a Highly Mobile MAGUK in Spines. J. Neurosci. 30, 4757–4766 (2010).

48. Metzbower, S. R., Levy, A. D., Dharmasri, P. A., Anderson, M. C. & Blanpied, T. A. Distinct SAP102 and PSD-95 Nano-organization Defines Multiple Types of Synaptic Scaffold Protein Domains at Single Synapses. J. Neurosci. 44, e1715232024 (2024).

49. Schamber, P. et al. SynAPSeg: A novel dataset and image analysis framework for deep learning-based synapse detection and quantification. bioRxiv 2026.03.12.711395 (2026) doi:10.64898/2026.03.12.711395.

50. Li, B. et al. Neuronal Inactivity Co-opts LTP Machinery to Drive Potassium Channel Splicing and Homeostatic Spike Widening. Cell 181, 1547–1565.e15 (2020).

51. Alkaas, A. et al. Synaptic PSD-95 biology: from localization and interactors to N-terminus function. Journal of Neurophysiology 134, 1588–1606 (2025).

52. Taylor, N. O., Wei, M.-T., Stone, H. A. & Brangwynne, C. P. Quantifying Dynamics in Phase-Separated Condensates Using Fluorescence Recovery after Photobleaching. Biophysical Journal 117, 1285–1300 (2019).

53. Morris, K. et al. Sequential replacement of PSD95 subunits in postsynaptic supercomplexes is slowest in the cortex. eLife 13, RP99303 (2024).

54. Kerr, J. M. & Blanpied, T. A. Subsynaptic AMPA Receptor Distribution Is Acutely Regulated by Actin-Driven Reorganization of the Postsynaptic Density. J. Neurosci. 32, 658–673 (2012).

55. Araki, Y. et al. SynGAP isoforms differentially regulate synaptic plasticity and dendritic development. eLife 9, e56273 (2020).

56. Tatavarty, V., Sun, Q. & Turrigiano, G. G. How to Scale Down Postsynaptic Strength. Journal of Neuroscience 33, 13179–13189 (2013).

57. Balzarotti, F. et al. Nanometer resolution imaging and tracking of fluorescent molecules with minimal photon fluxes. Science 355, 606–612 (2017).

58. Jeong, J. et al. PSD-95 binding dynamically regulates NLGN1 trafficking and function. Proc. Natl. Acad. Sci. U.S.A. 116, 12035–12044 (2019).

59. Futai, K. et al. Retrograde modulation of presynaptic release probability through signaling mediated by PSD-95–neuroligin. Nat Neurosci 10, 186–195 (2007).

60. MacGillavry, H. D., Song, Y., Raghavachari, S. & Blanpied, T. A. Nanoscale Scaffolding Domains within the Postsynaptic Density Concentrate Synaptic AMPA Receptors. Neuron 78, 615–622 (2013).

61. Böhme, M. A. et al. Active zone scaffolds differentially accumulate Unc13 isoforms to tune Ca2+ channel–vesicle coupling. Nat Neurosci 19, 1311–1320 (2016).

62. Reddy-Alla, S. et al. Stable Positioning of Unc13 Restricts Synaptic Vesicle Fusion to Defined Release Sites to Promote Synchronous Neurotransmission. Neuron 95, 1350–1364.e12 (2017).

63. Sakamoto, H. et al. Synaptic weight set by Munc13-1 supramolecular assemblies. Nat Neurosci 21, 41–49 (2018).

