## Supplemental material for "Btbd11 regulates glutamatergic synapse organization in GABAergic inhibitory interneurons"

### Figures and Tables

**Supplemental Figure 1.**
A) Btbd11 and PSD-95 predictive modeling. Predicted Aligned Error (PAE) plots of the interaction between Btbd11 and PSD-95, and Btbd11 and PSD-95-PDZ2Δ. Red box in the Btbd11-Psd-95 PAE plot represents binding site.

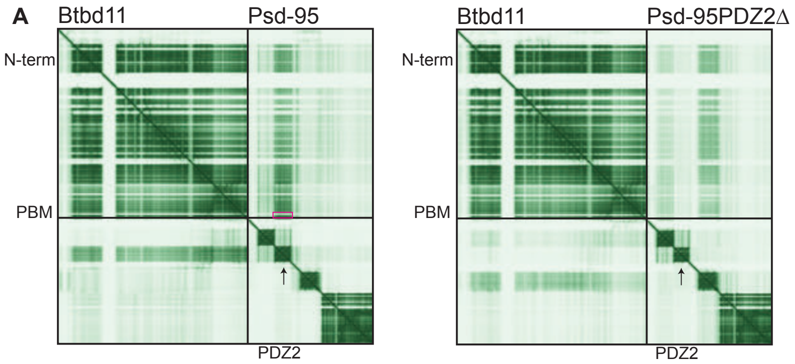

**Supplemental Figure 2.**
A) Individual channel breakdown of confocal images from Figure 2A. Scale bar = 10µm. B) FuzDrop predictive residue-promoting probability of Btbd11. C) Representative coalescence over time of multiple droplets containing PSD-95, TARPγ2-CT and Btbd11, a hallmark property of LLPS. Scale bar = 2µm. D) FuzDrop predictive residue-promoting probability of Btbd11ΔN. E) Representative confocal image of protein aggregate with azurite-Psd-95, GFP-TARPγ2-CT, and mCherry-Btbd11ΔN in HEK cells. Scale bar = 5µm. F) Representative confocal images show photobleaching of TARPγ2-CT in droplets containing PSD-95 and Btbd11ΔN over time. Scale bar = 2µm. G) One phase-association best-fit lines of TARPγ2-CT recovery in droplets containing PSD-95 and Btbd11ΔN show reduced recovery compared to PSD-95 and full-length Btbd11. H) Comparison of TARPγ2-CT recovery in droplets containing Psd-95 and Btbd11 or Btbd11ΔN. Unpaired t-test p = 0.0007, 4-8 replicates from 3 batches of cells. Error bars represent SEM.

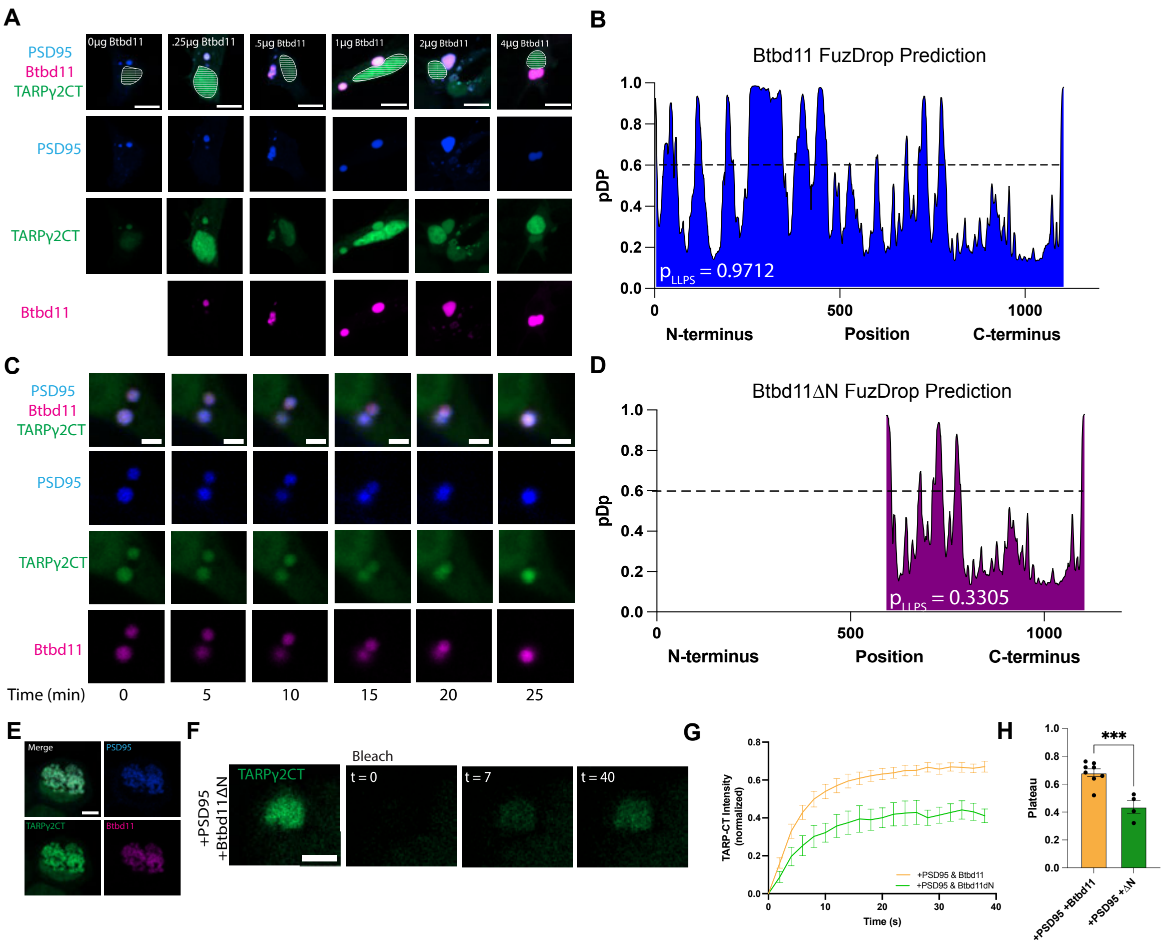

**Supplemental Figure 3.**
A) Raw STED images of *en face* synapse from main figure. Scale bar = 0.01μm. B) Correlation between synapse size and nanocluster number for Psd-95 and Btbd11. Btbd11 Pearson r=0.796, p=3.81e-217, Psd-95 Pearson r = 0.72, p=2.24e-158, n = 948 synapses from 3 independent batches of neurons. Shading represents SEM. C) Comparison of mean intersection of union (IoU) score of Psd-95 and Btbd11 per synapse. Mann-Whitney test p = .1545, Psd-95 n = 545 colocalized NCs, Btbd11 n = 482 colocalized NCs across 3 independent batches of cells. Error bars represent SEM. D) Correlation between Psd-95 and Btbd11 nanocluster size. Pearson r=0.14, p = p = 2.97 e-05, n = 948 synapses from 3 independent batches of neurons. Shading represents SEM. E) Comparison of mean Psd-95 and Btbd11 nanocluster size. Mann-Whitney test p < 0.0001, Psd-95 n = 3028 synapses, Btbd11 n = 2686 synapses from 3 independent batches of neurons. Error bars represent SEM. F) Raw STED images of side synapse from main figure. Scale bar = 0.01μm. G) Quantification of normalized maximum peak distance relative to Psd-95 for Btbd11 and Bassoon, taken from line scan data. Friedman test p < 0.0001, n = 93 synapses. Error bars represent SEM.

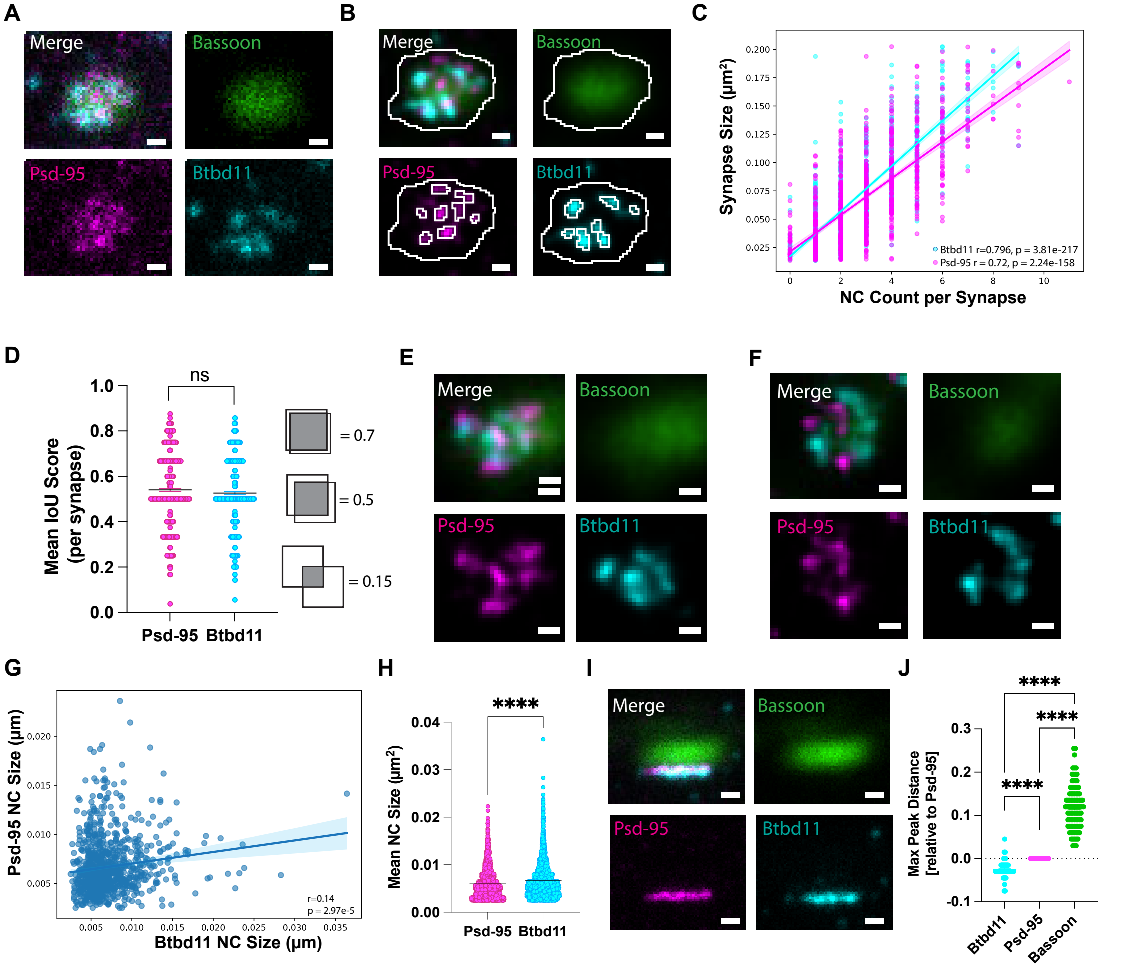

**Supplemental Figure 4.**

A) AlphaFold3 modeling of Btbd11 and SAP102 interaction predicts interaction through Btbd11 PBM to Psd-95 PDZ2. B) A representative immunoblot of GST-Btbd11 and mCherry-SAP102 shows detection of SAP102. SAP102 only was included as negative binding control.

C) Representative confocal imaging of colocalized GFP-Btbd11 and mCherry-SAP102 in HEK cells. Scale bar = 10μm. D) Representative immunoblot of GST-Btbd11ΔPBM and mCherry-SAP102 shows no detection of SAP102. SAP102 only was included as negative binding control. E) Quantification of mCherry-SAP102 pulled down by GST-Btbd11 (WT) or GST-Btbd11ΔPBM (ΔPBM) normalized signal shows significantly decreased detection of SAP102 with mutant (one-sample t-test $p < 0.0001$, $n = 3$ replicates). F) Representative immunoblot of GST-Btbd11ΔN and mCherry-SAP102 shows detection of SAP102. SAP102 only was included as negative binding control. G) Quantification of mCherry-SAP102 pulled down by GST-Btbd11 or GST-Btbd11ΔN shows no change in SAP102 detection, $n = 3$ replicates. H) Representative live-cell imaging of primary cultured rat hippocampal neurons transfected with mDlx-azurite and mCherry-SAP102 alone, with GFP-Btbd11, or with GFP-Btbd11ΔN. Lower panels display example individual bleached puncta and time-lapse recovery. Scale bar = 5μm. I) Normalized mCherry-SAP102 recovery with one-phase association best-fit line. J) GFP-Btbd11 overexpression significantly decreased SAP102 puncta plateau compared to Control and ΔN. One-way ANOVA $F = 6.109$, $p = 0.0048$, $n = 14\text{-}15$ cells from 5 independent cultures. K) Representative blot evaluating the interaction between GST-Btbd11 and myc-GluA1 following GST pulldown shows no detection of GluA1, $n = 3$ replicates. L) Representative confocal images of reduced GFP-Btbd11 expression following co-expression with Btbd11 CRISPR knockdown plasmid in HEK cells. Scale bar = 30μm.

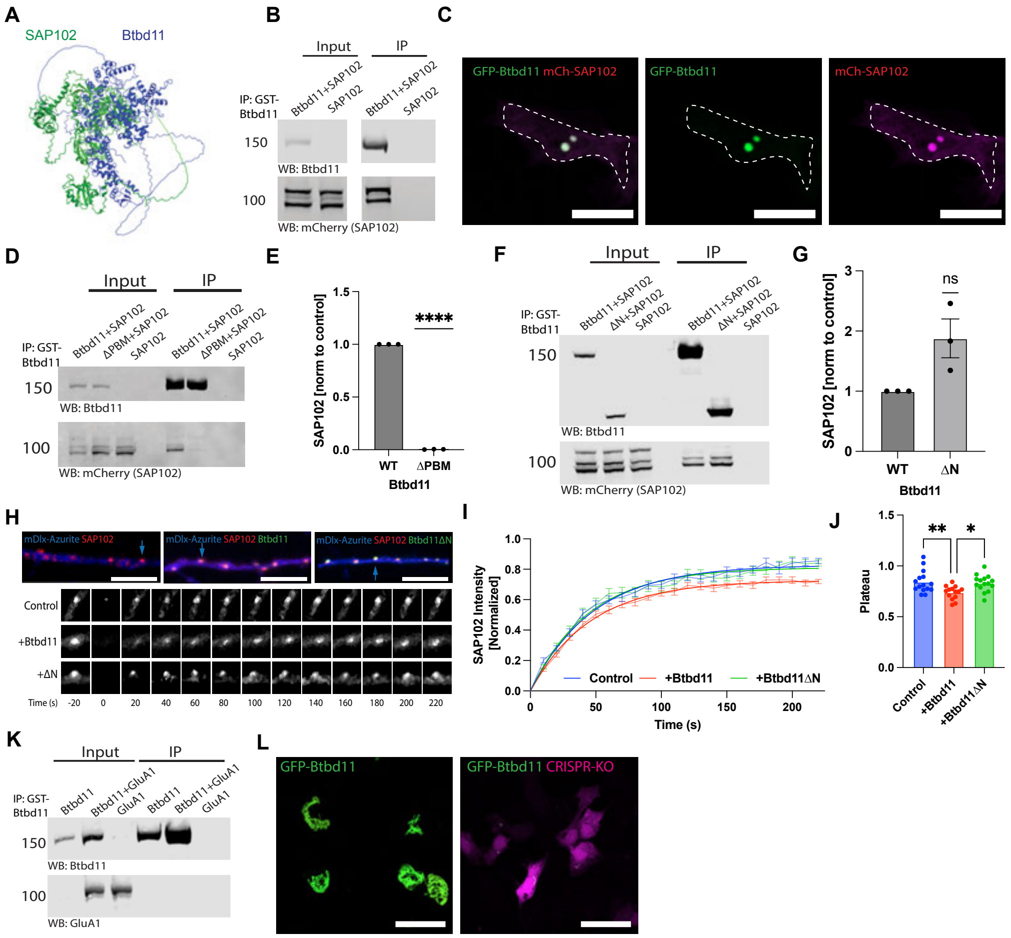

**Supplemental Figure 5.**

A) Representative synapse (upper panel) puncta detections (lower panels) for Psd-95 and VGlut1. Scale bars represent 5µm. B) Representative synapse (upper panel) puncta detections (lower panels) for TARPγ2 and VGlut1. Scale bars represent 5µm. C) Representative synapse (upper panel) puncta detections (lower panels) for GluA1 and VGlut1. Scale bars represent 5µm.

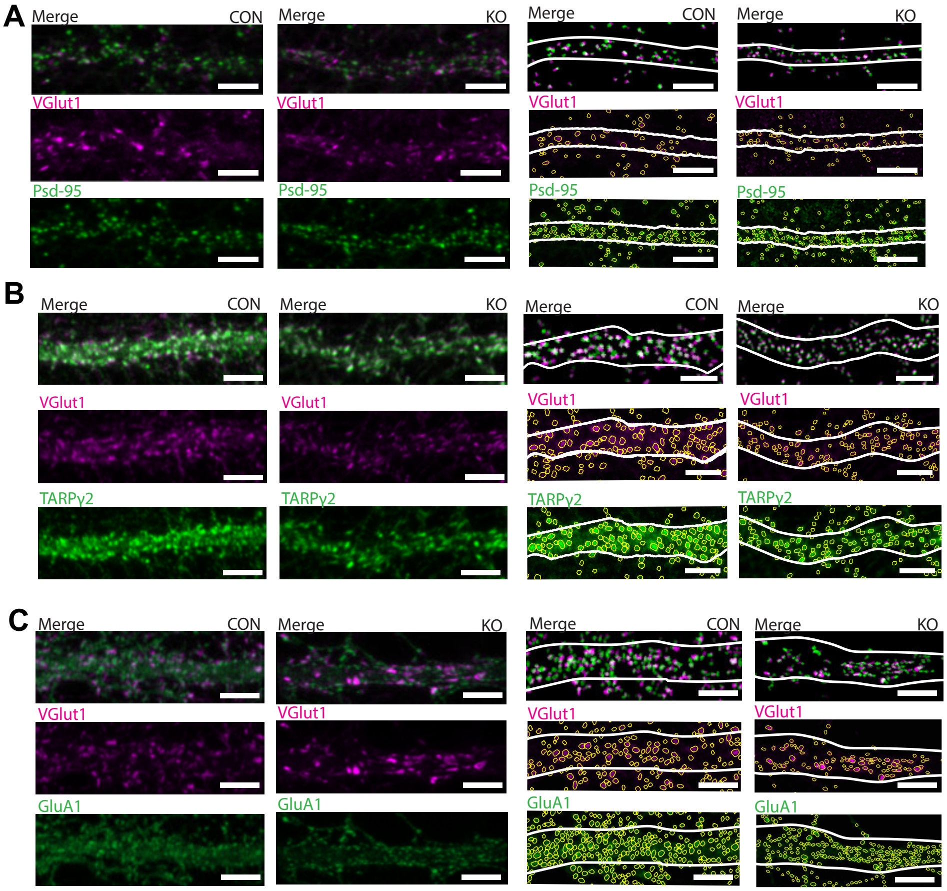

**Supplemental Figure 6.**

A) Comparison of average number of calcium events at baseline conditions in control and knockout neurons, unpaired t-test p = 0.0114, n = 13-14 neurons per condition from 2 batches of cells. Error bars represent SEM. B) Comparison of average number of calcium events in control at baseline and following AP5 and application, unpaired t-test p = 0.3905, n = 13-20 neurons per condition from 2-3 batches of cells. Error bars represent SEM. C) Comparison of average number of calcium events in knockout at baseline and following AP5 and application, Welch's t-test p = 0.1441, n = 14-20 neurons per condition from 2-3 batches of cells. Error bars represent SEM. D) Comparison of average number of calcium events in control and knockout neurons following AP5 and DNQX application, unpaired t-test p = 0.681, n = 13 neurons per condition from 2 batches of cells. Error bars represent SEM.

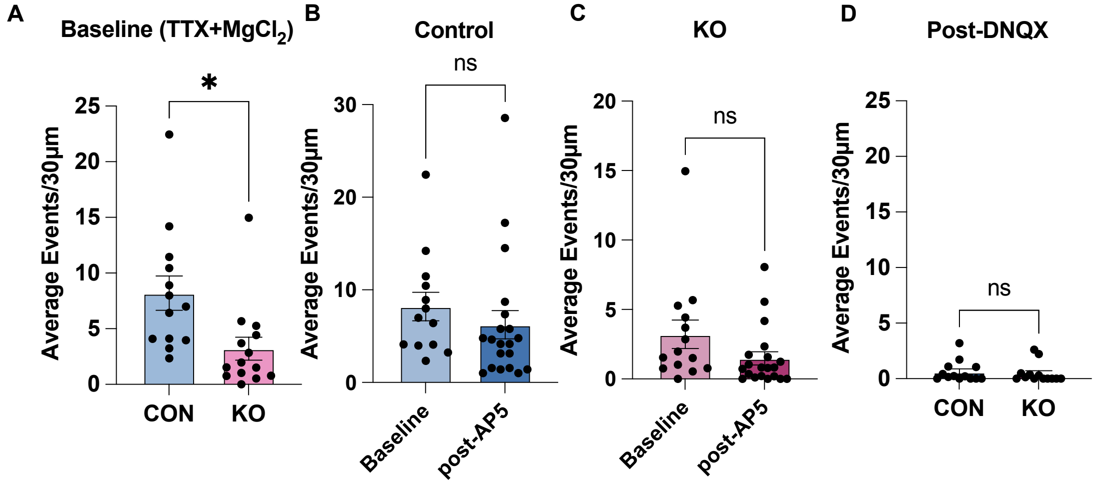

**Supplemental Figure 7.**

A) Comparison of synaptic Psd-95 area, averaged per cell. Unpaired t-test, $p = 0.0021$, $n = 18$ neurons from 3 batches of cells. Error bars represent SEM. B) Comparison of synaptic Psd-95 count, averaged per cell. Unpaired t-test, $p = 0.0356$, $n = 18$ neurons from 3 batches of cells. Error bars represent SEM. C) Correlation between synapse size and number of nanoclusters per synapse for control and Btbd11 KO. Control $r = 0.862$, KO $r = 0.849$. Control = 2327 synapses, KO = 1833 synapses from 3 batches of cells. Shading represents SEM.

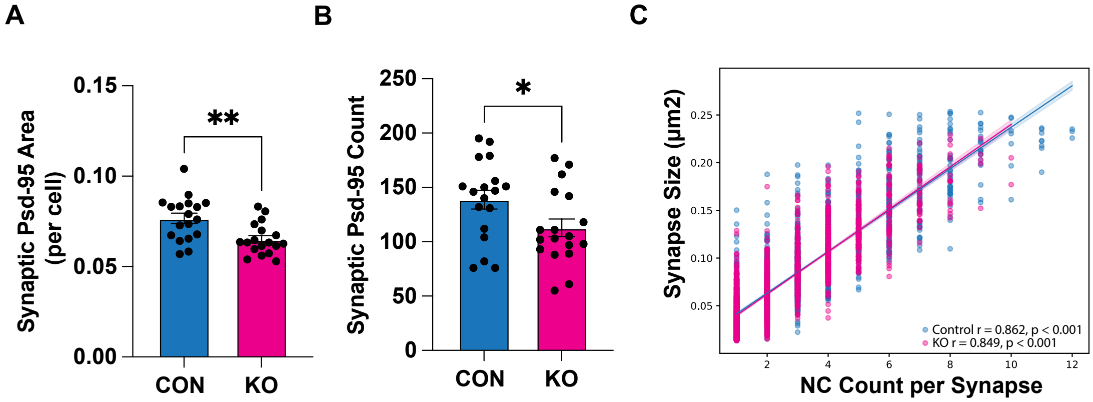

**Table 1.** Summary Stats Table.

| Figure | Test | Test Statistic | P-value | N number |
|---|---|---|---|---|
| Figure 1C | One-way ANOVA with Dunnett's multiple comparison | F(4,12)=129.9 | < 0.0001 | N = 3 per group |
| | Dunnett's multiple comparison | WT vs. Δ1,2 Mean Diff = 0.9706 | <0.0001 | |
| | Dunnett's multiple comparison | WT vs. Δ1 Mean Diff = 0.5723 | <0.0001 | |
| | Dunnett's multiple comparison | WT vs. Δ2 Mean Diff = 0.9589 | <0.0001 | |
| Figure 1E | One-way ANOVA with Dunnett's multiple comparison | F(4,108) | <0.0001 | N = 108 droplets |
| | Dunnett's multiple comparison | WT vs. Δ1,2 Mean Diff = 9.580 | <0.0001 | WT n = 40 droplets, Δ1,2 n = 23 droplets |
| | Dunnett's multiple comparison | WT vs. Δ1 Mean Diff = 0 5.441 | <0.0001 | WT n = 40 droplets, Δ1 n = 37 droplets |
| | Dunnett's multiple comparison | WT vs. Δ2 Mean Diff = 9.083 | <0.0001 | WT n = 40 droplets, Δ2 n = 8 droplets |
| Figure 1F | One-way ANOVA with Kruskal-Wallis test | F(4,43) | <0.0001 | N = 10-11 cells per condition |
| | Dunnett's multiple comparison | WT vs. Δ1,2 Mean Diff = 9.580 | <0.0001 | N = 11 cells |
| | Dunnett's multiple comparison | WT vs. Δ1 Mean Diff = 0 5.441 | >0.9999 | N = 11 cells |
| | Dunnett's multiple comparison | WT vs. Δ2 Mean Diff = 9.083 | <0.0001 | WT n = 11, Δ2 = 10 cells |
| Figure 1I | One-sample t-test (Psd-95) | t = 0.9243, df = 5, discrepancy = 0.02311 | P = 0.8731 | N = 6 |
| Figure 1J | One-sample t-test (TARP) | t = 3.023, df = 4, discrepancy = -.5039 | P = 0.0391 | N = 5 |
| Figure 2B | Btbd11 | F (4, 47) = 2.232 | P=0.0798 | 9-11 cells per condition |
| | (TARP) One-way ANOVA | F (5, 53) = 0.6257 | P=0.6808 | N = 8-10 cells per condition |
| | (PSD95) One-way ANOVA | F (5, 59) = 7.805 | P<0.0001 | N = 10-11 cells per condition |
| | (Droplet Size) One-way ANOVA with Kruskal-Wallis test | Kruskal-Wallis statistic = 63.82 | <0.0001 | Psd-95 and TARP only n = 44 droplets, + 0.25μg Btbd11 n = 46 droplets, |

| | | | | |
|---|---|---|---|---|
| | | | | +0.5μg Btbd11 n = 49 droplets, +1μg Btbd11 n = 61 droplets, +2μg Btbd11 n = 54 droplets, +4μg Btbd11 n = 55 droplets |
| Figure 2C (IP TARP signal) | One-way ANOVA with Kruskal-Wallis test | Kruskal-Wallis statistic = 1.223 | p = 0.8742 | N = 4 replicates |
| Figure 2D (Droplet size) | Welch ANOVA | W (DFn, DFd) 9.053 (2.000, 14.67) | p = 0.0027 | N = 7-12 droplets per condition from 3 batches of cells |
| Figure 2G (Plateau) | Unpaired t-test | t=0.7912, df=13.08 | 0.4420 | N = 8 droplets per condition from 3 batches of cells |
| Figure 2H (droplet size) | Welch's t-test | t=0.7765, df=8.764 | p = 0.4579 | N = 8 droplets per condition from 3 batches of cells |
| Supp 2H | Unpaired t-test | t=4.840, df=10 mean difference = -0.2453 ± 0.05068 | P = 0.0007 | +Psd-95+ Btbd11 n = 8 droplets, +Psd-95+ Btbd11 n 4 aggregates |
| Figure 3C (Intensity correlation) | Pearson Correlation | r = 0.36 | p = 3.133e-158 | N = 5073 synapses |
| Figure 3E (% NC coloc) | Mann-Whitney test | Mann-Whitney U = 456837, difference = 0.06667 | P = 0.2037 | Psd-95 n = 953 synapses, Btbd11 n = 991 synapses |
| Figure 3F (NC count correlation) | Pearson Correlation | r=0.67 | P = 4.32 e-123 | n = 948 synapses |
| Figure 3I (Feret diameter) | Paired Wilcoxon test | Median of differences = -.032 | P < 0.0001 | N = 120 synapses |
| Supp 3B (Synapse-NC count correlation) | Pearson correlation | Psd-95 r = 0.72 Btbd11 r = r=0.796 | Psd-95 p = 2.24e-158 Btbd11 p = 3.81e-217 | n = 948 synapses |
| Supp 3C (IOU score) | Mann-Whitney test | Mann-Whitney U = 124748, difference = 0 | P = 0.1545 | Psd-95 n = 545, Btbd11 n = 482 synapses |

| | | | | |
|---|---|---|---|---|
| Supp 3D (NC size correlation) | Pearson Correlation | r = 0.14 | p = 2.97 e-05 | n = 948 synapses |
| Supp 3E (NC size) | Mann-Whitney test | Mann Whitney U = 3649123, Psd-95 mean = 0.006003, Btbd11 mean = 0.006732 | P < 0.0001 | Psd-95 n = 3028 NCs, Btbd11 n = 2686 NCs |
| Supp 3G (Peak distance) | Friedman test | Friedman statistic = 172.2 | P < 0.0001 | N = 92 synapses |
| Figure 4C (Btbd11 FRAP) | Welch's t-test | t=6.337, df=6.846 mean diff = -0.6285 ± 0.09919 | p = 0.0004 | Btbd11 n = 7 cells, ΔN n = 6 cells from 6 batches |
| Figure 4F (TARP FRAP) | One-way ANOVA with Dunnett's multiple comparisons | F (2, 24) = 5.776 | P=0.0090 | N = 9-10 neurons per condition from 5 batches |
| | Dunnett's multiple comparisons | Control vs +Btbd11, mean diff 0.1326 | P = 0.0172 | Control n = 10 neurons, +Btbd11 n = 9 neurons |
| | Dunnett's multiple comparisons | +Btbd11 vs +ΔN, mean diff -0.1364 | P = 0.0207 | N = 9 neurons for both conditions |
| | Dunnett's multiple comparisons | Control vs +ΔN, mean diff = -0.003840 | P = 0.9962 | Control n = 10 neurons, +ΔN n = 9 neurons |
| Figure 4I (GluA1 FRAP) | One-way ANOVA with Dunnett's multiple comparisons | F (2, 22) = 15.01 | P < 0.0001 | N = 7-9 neurons per condition from 4 batches |
| | Dunnett's multiple comparisons | Control vs +Btbd11, mean diff -0.1792 | P = 0.0001 | N = 9 neurons for both conditions |
| | Dunnett's multiple comparisons | +Btbd11 vs +ΔN, mean diff 0.1572 | P = 0.0011 | +Btbd11 n = 9 neurons, +ΔN n = 9 neurons |
| | Dunnett's multiple comparisons | Control vs +ΔN, mean diff = -0.02193 | P = 0.8309 | Control n = 9 neurons, +ΔN n = 7 neurons |
| Figure 4L (GluA1 FRAP, Btbd11 KO) | Unpaired t-test | t=3.448, df=13, mean diff = -0.2247 | p = 0.0043 | Control n = 7 neurons, KO n = 8 neurons from batches |
| Supp 4E (ΔPBM-SAP102 IP) | One-sample t-test | t=1125, df=2 discrepancy = -0.9960 | P < 0.0001 | 3 replicates |

| | | | | |
|---|---|---|---|---|
| Supp 4G (ΔN-SAP102 IP) | One sample t-test | t=2.729, df=2, discrepancy = 0.8790 | P = 0.1122 | 3 replicates |
| Supp 4J (SAP102 FRAP) | One-way ANOVA with Dunnett's multiple comparisons | F (2, 41) = 6.109 | P=0.0048 | N = 14-15 neurons from 5 batches |
| | Dunnett's multiple comparisons | Control vs +Btbd11, mean diff = 0.1043 | P = 0.0071 | Control n = 15 neurons, +Btbd11 n = 14 neurons |
| | Dunnett's multiple comparisons | +Btbd11 vs +ΔN, mean diff = -0.09230 | P = 0.0187 | +Btbd11 n = 14 neurons, +ΔN n = 15 neurons |
| | Dunnett's multiple comparisons | Control vs +ΔN, mean diff = 0.01204 | P = 0.9247 | N = 15 neurons for both conditions |
| Figure 5C (Psd-95 Area) | Unpaired t-test with Welch's correction | t=2.348, df=37.49 mean diff= -0.03608 ± 0.01536 | P = 0.0242 | N = 25 neurons from 4 batches per condition |
| (Psd-95 Density) | Unpaired t-test | t=2.402, df=48, mean diff = -0.09404 ± 0.03915 | P = 0.0202 | |
| Figure 5F (TARPγ2 Intensity) | Unpaired t-test | t=2.379, df=47 mean diff = -0.2448 ± 0.1029 | P = 0.0215 | Control n = 24 neurons, KO n = 25 neurons from 4 batches |
| (TARPγ2 Density) | Unpaired t-test | t=4.230, df=46 mean diff = -0.1249 ± 0.02953 | P = 0.0001 | |
| Figure 5I (GluA1 intensity) | Unpaired t-test | t=2.956, df=46, mean diff = -0.1334 ± 0.04514 | 0.0049 | N = 24 neurons from 4 batches per condition |
| Figure 5I (GluA1 density) | Unpaired t-test with Welch's correction | t=3.617, df=37.28, mean diff = -0.2457 ± 0.06793 | P = 0.0009 | N = 24 neurons from 4 batches per condition |
| Figure 6E (Peaks) | Unpaired t-test with Welch's correction | t=6.459, df= 488.43, mean diff =4.695 | P < 0.0001 | Control = 612 events, KO = 184 events |
| Figure 6F (ΔF/F) | Mann-Whitney test | Mann-Whitney U = 111, difference = 0.09472 | P = 0.0262 | Control n = 20 neurons, KO n = 19 neurons |
| Figure 6G (Frequency) | Mann-Whitney test | Mann-Whitney test U = 38, difference = 5.928 | P < 0.0001 | N = 20 neurons per condition |
| Supp 6A (Baseline) | Unpaired t-test | t=2.733, df=25, mean diff = -4.988 ± 1.826 | P = 0.0114 | Control n = 13 neurons, KO n = 14 neurons |

| | | | | |
|---|---|---|---|---|
| Supp 6B (DNQX) | Unpaired t-test | t=0.4162, df=24, mean diff = 0.1495 ± 0.3592 | P = 0.681 | N = 13 neurons per condition |
| Supp 6C | Unpaired t-test | t=0.8710, df=31, mean diff = 1.967 | P = 0.3905 | Baseline n = 13 neurons, post-AP5 n = 20 neurons |
| Supp 6D | Unpaired t-test with Welch's correction | t=1.525, df=18.57, mean diff = -1.724 ± 1.130 | P = 0.1441 | Baseline n = 14 neurons, post-AP5 n = 20 neurons |
| Figure 7C (NC/synapse, cell avg) | Unpaired t-test | t=2.997, df=34 mean diff = -0.3849 ± 0.1284 | P = 0.0051 | N = 18 neurons from 3 batches per condition |
| Figure 7D (NC/synapse, all) | Mann-Whitney test | Mann-Whitney U = 1842914, median difference = 0 (median = 2 for each condition) | P < 0.0001 | Control n = 2363 synapses, KO n = 1863 synapses |
| Figure 7E (NC Area, cell avg) | Unpaired t-test | t=3.039, df=34 mean diff = 0.0002734 ± 8.996e-005 | P = 0.0045 | N = 18 neurons from 3 batches per condition |
| Figure 7F (NC Area, all) | Mann-Whitney test | Mann-Whitney U = 17430825, difference = 0 (median = 0.003 for each condition) | P < 0.0001 | Control n = 7390 NCs, KO n = 4957 NCs |
| Figure 7J Psd-95 depth, cell avg) | Unpaired t-test with Welch's correction | t=0.8809, df=25.19, mean diff = -0.01036 ± 0.01176 | P = 0.3867 | Control n = 18 neurons, KO n = 17 neurons from 3 batches per condition |
| Figure 7K (Psd-95 depth, all) | Mann-Whitney test | Mann-Whitney U = 8604, difference = 0.01050 | P = 0.1539 | Control n = 162 synapses, KO = 118 synapses |
| Figure 7L (Psd-95 width, cell avg) | Unpaired t-test | t=3.349, df=33, mean diff = -0.06562 ± 0.01959 | P = 0.002 | Control n = 18 neurons, KO n = 17 neurons from 3 batches per condition |
| Figure 7M (Psd-95 width, all) | Mann-Whitney test | Mann-Whitney U = 7060, difference = 0.067 | P <0.0001 | Control n = 163 synapses, KO = 124 synapses |
| Supp 7A (Synaptic Psd-95 Area) | Unpaired t-test | t=3.331, df=34, mean diff = 0.01163 ± 0.003493 | P = 0.0021 | 18 neurons from 3 batches of cells |
| Supp 7B (Synaptic Psd-95 Count) | Unpaired t-test | t=2.188, df=34, mean diff = 26.06 ± 11.91 | P = 0.0356 | 18 neurons from 3 batches of cells |

| | | | | |
|---|---|---|---|---|
| Supp 7C (NC Count-Synapse size correlation) | Pearson Correlation | Control r = 0.862, KO r = 0.849 | P < 0.0001 for both | Control = 2327 synapses, KO = 1833 synapses |
